# Robust self-supervised denoising of voltage imaging data using CellMincer

**DOI:** 10.1101/2024.04.12.589298

**Authors:** Brice Wang, Tianle Ma, Theresa Chen, Trinh Nguyen, Ethan Crouse, Stephen J. Fleming, Alison S. Walker, Vera Valakh, Ralda Nehme, Evan W. Miller, Samouil L. Farhi, Mehrtash Babadi

## Abstract

Voltage imaging enables high-throughput investigation of neuronal activity, yet its utility is often constrained by a low signal-to-noise ratio (SNR). Conventional denoising algorithms, such as those based on matrix factorization, impose limiting assumptions about the noise process and the spatiotemporal structure of the signal. While deep learning based denoising techniques offer greater adaptability, existing approaches fail to fully exploit the fast temporal dynamics and unique short- and long-range dependencies within voltage imaging datasets. Here, we introduce CellMincer, a novel self-supervised deep learning method designed specifically for denoising voltage imaging datasets. CellMincer operates on the principle of masking and predicting sparse sets of pixels across short temporal windows and conditions the denoiser on precomputed spatiotemporal auto-correlations to effectively model long-range dependencies without the need for large temporal denoising contexts. We develop and utilize a physics-based simulation framework to generate realistic datasets for rigorous hyperparameter optimization and ablation studies, highlighting the key role of conditioning the denoiser on precomputed spatiotemporal auto-correlations to achieve 3-fold further reduction in noise. Comprehensive benchmarking on both simulated and real voltage imaging datasets, including those with paired patch-clamp electrophysiology (EP) as ground truth, demonstrates CellMincer’s state-of-the-art performance. It achieves substantial noise reduction across the entire frequency spectrum, enhanced detection of subthreshold events, and superior cross-correlation with ground-truth EP recordings. Finally, we demonstrate how CellMincer’s addition to a typical voltage imaging data analysis workflow improves neuronal segmentation, peak detection, and ultimately leads to significantly enhanced separation of functional phenotypes.

## 1 Introduction

Voltage imaging utilizes fluorescent reporters, either small-molecule dyes or genetically encoded proteins, to measure the membrane potential of electrically active cells. Compared to traditional patch-clamp electrophysiology (EP), voltage imaging offers higher throughput and is less invasive. This technique has been used to monitor neuronal electrical activity during behavioral assays *in vivo* [1], as well as to characterize the functional effects of pharmacological and genetic perturbations in primary and iPSC-derived mammalian neurons *in vitro* [2, 3]. The increased throughput, control, and flexibility of voltage imaging have enabled significant advances in our understanding of biology. Recent developments have improved the brightness of both voltage-sensitive dyes [4] and heterologously expressed voltage-sensitive proteins [5]. However, the achievable signal-to-noise ratio (SNR) remains limited compared to conventional patch-clamp techniques due to factors such as dye quantum yield, short exposure times (*<*2 ms) needed to capture neuronal action potentials, and constraints on excitation intensity to prevent sample damage.

Limitations on SNR have two practical effects. First, small-magnitude electrical events of interest, such as subthreshold post-synaptic potentials, could be lost in the background temporal noise. Previous work powered by tracking the timing of action potential firing between neurons provided some insight to how neurons might wire together in small circuits. However, the site of inter-neuronal communication is the synapse. A measure of synaptic connectivity, demonstrated as subthreshold activity measured at the soma, would provide a greater understanding of how neuronal circuits are formed and how synaptic connections are modified during different forms of plasticity. Second, cells expressing relatively low amounts of fluorescent reporters can be lost in a comparatively high autofluorescent background, lowering the effective throughput of voltage imaging.

These technical challenges have motivated the development of data denoising algorithms to computationally enhance the SNR and enable the recovery of obscured and subtle fluorescent signals. Matrix factorization is an effective class of algorithms for fluorescence image denoising [6] [7], as the sparse and static signal sources (e.g. neurites) in these imaging assays create an ideal setting for approximating entire fluorescence recordings as low-rank decompositions. Principal component analysis (PCA), non-negative matrix factorization (NMF), and penalized matrix decomposition (PMD) [8] are popular implementations of this concept. These approaches, while being highly efficient and effective at data denoising, suffer from a number of caveats. These include: (1) implicit parametric assumptions on the nature of the noise that are theoretical approximation of the actual complex data generating process; (2) usage of spatiotemporal regularizations to encourage robustness and model identifiability, such as total variation penalty or temporal continuity, that are often violated (e.g. spike events, spatially heterogeneous expression of the fluorescent reporter); (3) making strong assumptions about the background fluorescence component to allow their approximate subtraction as a simple data preprocessing step. These modeling assumptions, while laying a strong foundation, ultimately hamper the expressivity of conventional denoising algorithms.

We envision that the ideal denoising algorithm should minimize the explicit assumptions made about the noise process while maximizing the potential to learn the complex spatiotemporal relationships that govern the signal. Deep neural networks (DNNs), which have no theoretical limit to complexity, can in principle solve the issue of denoising model expressivity. However, deep learning denoising models pretrained on large datasets of natural images [9] are ill-equipped to operate in the low-SNR regime of fluorescence imaging [10], requiring a suitable model to be trained from scratch for this data domain. This immediately poses a challenge for supervised learning approaches which require clean images as a learning target, which are not available in voltage imaging. A powerful recent training strategy, called self-supervised denoising, circumvents the requirement of having clean ground truth data by exploiting a key property of many noise processes: by appropriately partitioning the raw data into compartments, and predicting one compartment from the other, it is often possible to eliminate predictors of noise while retaining the ability to predict the underlying signal. Noise2Noise (N2N) [11] and Noise2Self (N2S) [12] are prominent examples of self-supervised denoising techniques proposed for images. These methods have consistently been shown to produce state-of-the-art results, including in fluorescence imaging, even compared to counterparts that are trained on pairs of noisy and clean data [10]. In particular, the Noise2Self algorithm, which we use as a foundation to build upon here, operates on the following elegant and simple principle: suppose a sparse set of pixels are masked out from a noisy image, and a neural network is trained to predict the value of the sparsely masked pixels from the rest of the image, i.e. the majority of pixels. Assuming that the noise in the masked pixels is uncorrelated with the rest of the pixels, the optimal predictor can at best predict the noiseless *signal* component; in practice, it can excel at this task given the strong spatial correlations and redundancy in biological images. It follows that the optimal masked pixel predictor in turn behaves as an optimal pixel denoiser. Noisy data itself provides the needed evidence for teasing out the signal component, circumventing the need for clean training data.

One of the main challenges in extending self-supervised image denoising approaches to spatiotemporal data, such as voltage imaging recordings, is that these datasets contain thousands of frames, and that each frame is individually too signal-deficient to self-supervise its own denoising. At the same time, GPU hardware memory constraints and efficient training considerations prevent us from ingesting and processing entire voltage imaging movies with neural networks to exploit frame-to-frame correlations. The middle ground strategy adopted by several authors is to process the movie in overlapping and truncated *local denoising temporal contexts*, i.e. chunks of adjacent frames. For instance Li et al. [13] developed DeepCAD, a Noise2Self-like denoising method based on reconstructing whole masked frames from temporally downsampled movie chunks, and demonstrated its capacity to restore a high imaging SNR from low-SNR calcium imaging recordings. Lecoq et al. [14] developed DeepInterpolation, a Noise2Noise-like whole-frame interpolation-based deep learning method acting on small temporal windows which also allowed them to increase the SNR and retrieve a significantly higher fraction of neuronal segments from calcium imaging.

These existing approaches are sub-optimal for two important reasons. (1) As we will show in later sections, while such leave-frame-out approaches work remarkably well for calcium imaging, the denoising performance degrades strikingly when the same methods are applied to voltage imaging data (see Sec. 2.3). The much faster temporal dynamics of voltage imaging data compared to calcium imaging imply that each movie frame contains unique and valuable signal that cannot be entirely inferred from the adjacent frames. For instance, the evidence for a neuronal spike is most prominently present in a single frame. (2) Unless the local denoising context is impractically large and contains hundreds of movie frames, the neural network is incapable of estimating long-range pixel-to-pixel temporal correlations that are arguably key to effective signal extraction and noise removal. As we will show in later sections, explicitly precomputing and supplementing the short-context local denoiser with such information results in a striking boost in the denoising performance.

In this work, we introduce CellMincer, a self-supervised deep learning method specifically designed for denoising voltage imaging datasets based on the Noise2Self denoising framework. CellMincer introduces several key refinements over the currently existing self-supervised movie denoising methods to address the aforementioned caveats. The key methodological contributions of CellMincer include: (1) development of an efficient and expressive two-stage spatiotemporal data processing deep neural network architecture, comprising a frame-wise 2D U-Net module for spatial feature extraction, followed by a pixelwise 1D convolutional module for temporal data post-processing; (2) replacing the common task of whole-frame prediction with masking and predicting a sparse set of pixels from a small number of adjacent frames; this training methodology allows the denoiser to have access to the unique information contained in any individual frame as well as the supporting context in its neighboring frames; (3) precomputing spatiotemporal auto-correlations at multiple length scales, and providing such precomputed statistics as a *conditioner* to the denoiser neural network (that otherwise processes smaller spatiotemporal regions of the movie at a time); (4) developing and leveraging a physics-based simulation framework to generate highly realistic pairs of clean *ground truth* and noisy recording realization for hyperparameter optimization and performing ablation studies to tease apart the roles of various modeling choices in a controlled setting.

Using benchmarking experiments performed on simulated data and real voltage imaging data with paired patch-clamp EP recordings as a proxy for ground truth, we show that CellMincer yields state-of-the-art results as measured in terms of several practical metrics. These include a peak signal-to-noise ratio (PSNR) average gain of 24 dB compared to the raw data (an increase of 2 dB over the next best benchmarked method), a 14 dB reduction in high-frequency (*>*100 Hz) noise (a further reduction of 10.5 dB from the next best method), a 5-10 percentage point increase of *F*_1_-score in detecting sub-threshold events compared to the other algorithms and across all voltage magnitudes in the 1-10 mV range (in which the baseline *F*_1_-score ranges from 5-14%), and more than 20% increase in the cross-correlation between low-noise EP recordings and voltage imaging. A striking result from our ablation study is the pivotal role of conditioning the denoiser on precomputed global features, resulting in a nearly 5 dB (or approximately 3-fold) boost in average PSNR gain, as well as a highly-concentrated distribution of PSNR gain across all frames and electrical stimulation amplitudes (see Fig. 1e). Finally, to demonstrate the utility of CellMincer to real end-to-end biological hypothesis testing, we compare the voltage imaging of chronically tetrodotoxin-treated and unperturbed cultured hPSC-derived neurons, and demonstrate that CellMincer denoising enables reliable identification and segmentation of nearly 2-fold as many neurons as in the raw data, improved identification of spiking events, and ultimately significantly enhanced statistical separation between the two functional phenotypes.

**Figure 1.**
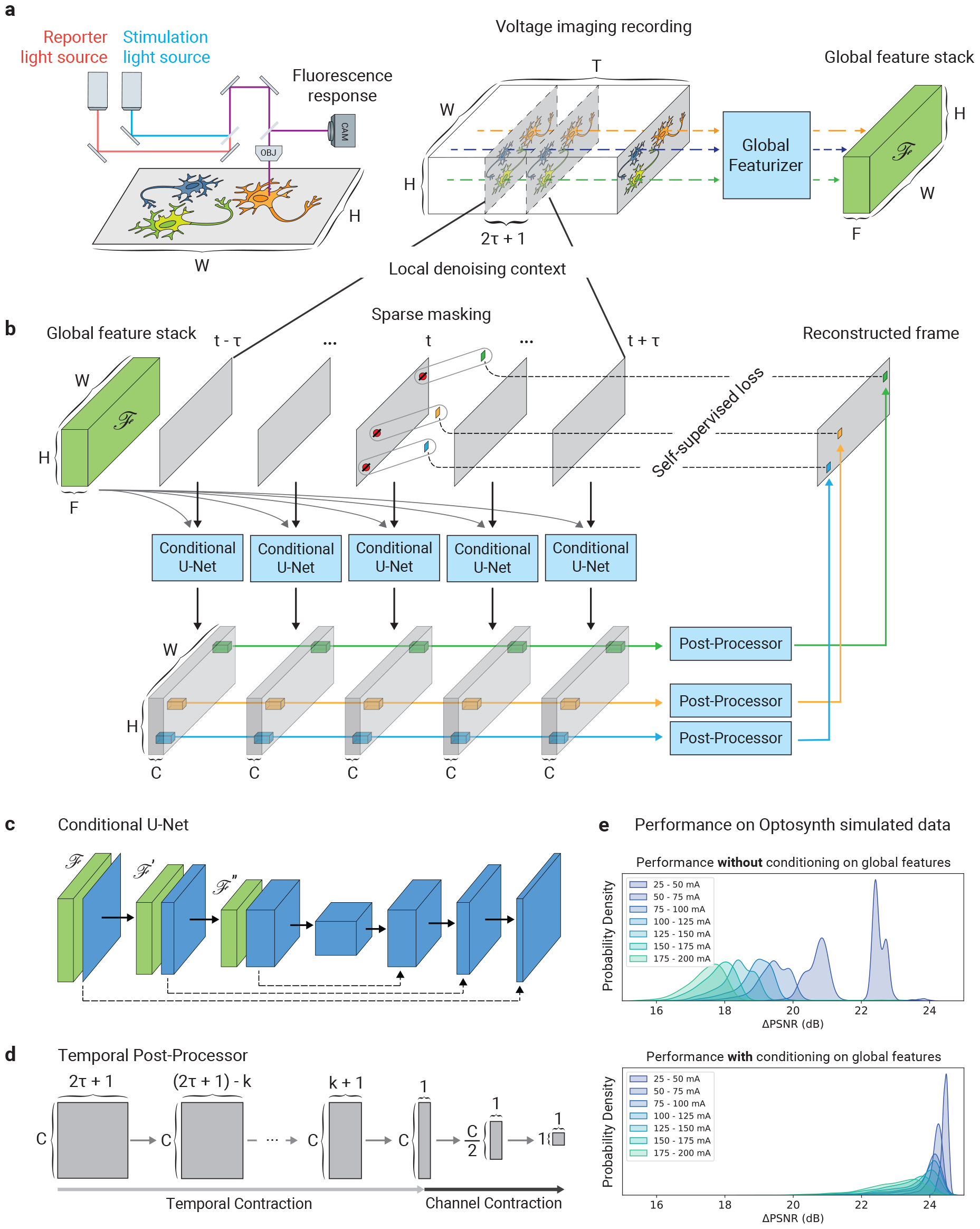
Overview of voltage imaging data and CellMincer denoising model. (a) A simplified schematic diagram of a typical optical voltage imaging experiment (left). The spatially resolved fluorescence response is recorded over time to produce a voltage imaging movie. A key component of CellMincer’s preprocessing pipeline is the computation of spatial summary statistics and various auto-correlations from the entire recording, which are concatenated into a stack of global features (right). (b) An overview of CellMincer’s deep learning architecture. (c) The conditional U-Net convolutional neural network (CNN). At each step in the contracting path, the precomputed global feature stack is spatially downsampled in parallel (*ℱ → ℱ*^*′*^ *→ ℱ*^*′′*^ *→* …) and concatenated to the intermediate spatial feature maps. (d) The temporal post-processor neural network. The sequence of pixel embeddings are convolved with a 1D kernel along the time dimension, producing a single vector of length *C*. A multilayer perceptron subsequently reduces this vector to a single value. (e) A comparison of model performance on simulated data before and after introducing global features as a U-Net conditioner. The distributions of PSNR gain are binned by stimulation amplitude. Using global features confers an average increase of 5 dB to the denoiser, roughly corresponding to a 3-fold noise reduction.

## 2 Results

### 2.1 CellMincer self-supervised denoising framework

The CellMincer denoising pipeline involves three stages: (1) data preprocessing and global feature extraction; (2) self-supervised pretraining of the denoising neural network; (3) inference of denoised movie. In the preprocessing step, we take a recording *X*(*t, x, y*) represented as a three-dimensional tensor with shape *T* (time) *× W* (width) *× H* (height). We treat each pixel as a separate time series of *T* samples, fit a low-order polynomial function to each, and thereby decompose the movie as a sum of *smooth trend* and *detrended residual* tensors. The trend tensor primarily represents the background fluorescence, whereas the residual detrended tensor represents a noisy measurement of the electrical activity. Going forward, we perform self-supervised denoising only over the detrended residual component, and add the smooth trend component back after the inference step. To set the stage for self-supervised pretraining, we calculate various summary statistics for each pixel, including temporal mean, temporal variance, and all bidirectionally lagged spatiotemporal auto-correlations with adjacent pixels. These statistics are computed separately for both the *slow moving average* and the *fast residual* components of the detrended movie, and at two different spatially downsampled resolutions to account for auto-correlations with longer spatial lags. Finally, we concatenate these precomputed statistics as a tensor *ℱ* of shape *F × W × H* to represent pixelwise global statistics, where *F* = 74 is the total number of computed statistics per pixel. Supplemental Sec. S.3 fully describes our preprocessing and feature extraction stage. This step is schematically referred to as *global featurizer* in Fig. 1a.

We present the denoising strategy we employ in CellMincer in two stages for clarity. First, we describe the architecture of the DNN that we purport to be capable to performing efficient denoising. Next, we describe the self-supervised training strategy *a la* Noise2Self that allows the denoiser to train without clean targets.

Our proposed denoising DNN takes as input a series of 2*τ* +1 consecutive frames, corresponding to time points *t − τ*, …, *t −* 1, *t, t* + 1, … *t* + *τ*, from the detrended movie and aims to predict a denoised reconstruction of the frame in the middle, at time point *t*. We refer to *τ* as the *temporal order*, and to 2*τ* + 1 as the *context size* of the local denoiser. Crucially, the DNN additionally takes the precomputed global feature stack *ℱ* as a conditioner to supplement the local denoiser with long-range spatiotemporal statistics. The architecture of the denoising DNN consists of a single-frame spatial feature extractor followed by a temporal post-processor (see Fig. 1b). The spatial component is implemented as a U-Net convolutional neural network (CNN) with a small but consequential modification: to condition the convolutional operations on *ℱ*, we concatenate an appropriately spatially downsampled copy of *ℱ* prior to each convolution block on the contracting path (see Fig. 1c). The conditional U-Net extracts deep, native-resolution *C*-channel single-frame embeddings from each of the 2*τ* + 1 consecutive frames (see Fig. 1b). The resulting embeddings are concatenated into a 4D tensor of shape (2*τ* + 1) *× C × W × H*:

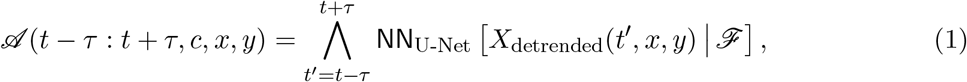

where ∧ denotes concatenation along the time dimension. This intermediate tensor is routed to the temporal post-processor, which consists of a series of temporal convolutional layers, reducing each set of pixel embeddings across all frames to a final output pixel. The output of the temporal post-processor represents a denoised reconstruction of the middle frame:

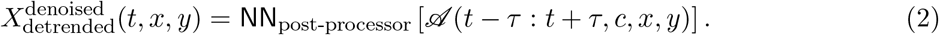

Note that the temporal post-processor treats pixels (*x, y*) as independent, only operating on time and U-Net feature channels (see Fig. 1d). This two-stage constrained network design enables efficient spatiotemporal data processing by logically compartmentalizing the flow of information; the U-Net facilitates information mixing across pixels within individual frames, while the post-processor convolves information across frames for individual pixels. Refer to Supplemental Sec. S.4 for architectural details.

We train the CellMincer denoiser in a self-supervised fashion as follows. At the beginning of each training iteration, pixels chosen at random in the frame at time *t* are replaced with Gaussian noise with pixel-specific in-distribution mean and variance before the frame is fed into the network:

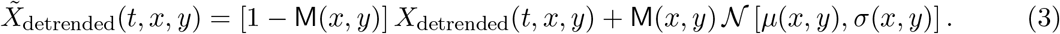

Here, M(*x, y*) is a binary mask containing a sparse number of ones, and *μ*(*x, y*) and *σ*(*x, y*) correspond to the temporal mean and standard deviation of the detrended movie at position (*x, y*). These masked pixels are then used as the training targets, where the *L*_*p*_ loss is computed between the network’s predicted values and their pre-masked values with the following loss function:

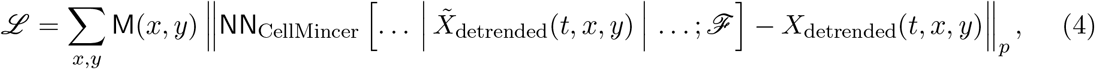

where NN_CellMincer_ = NN_post-processor_ ∘ NN_U-Net_ (see Fig. 1b). Here, … | and | … refer to the *τ* preceding and the *τ* following frames surrounding the frame at time *t*, respectively. As for a choice of pixel loss function, we have experimented with both *p* = 1 and 2 and found the latter to result in higher PSNR (see Supplemental Sec. S.6). Even though these pre-masked target values do not represent actual ground truth but noisy realizations, their noise contribution cannot be predicted by the neural network provided that the masking mechanism decorrelates the pixel noise between masked and unmasked compartments (*𝒥*-invariance, see Ref. [12]). In fluorescence imaging, the main source of noise is pixelwise Poisson-Gaussian noise, which is uncorrelated across pixels, allowing us to satisfy the *𝒥* -invariance condition as a matter of masking individual pixels. CellMincer’s implementation additionally permits the use of alternate masking mechanisms (e.g. inclusion of margin around each masked pixel) if needed to account for correlated noise. Crucially, since the identities of the masked pixels are not revealed to the denoiser (e.g. implicitly by using a special masked pixel value, as is typically done), the network is incentivised to denoise not only the sparse subset of masked pixels but the frame in its entirety. This training strategy allows CellMincer to operate very efficiently at inference time, when we feed the noisy detrended movie in (2*τ* + 1)-length overlapping sliding windows to the network and denoise each window’s entire middle frame. To avoid the typical practice of producing truncated results, we augment all model inputs with appropriate spatial padding at training and inference time, and we pad the beginning and end of the denoised movie with *τ* copies of its first and last frame respectively.

Our implementation of CellMincer can jointly train on many datasets across multiple GPUs to produce a highly generalizable model, but satisfactory results can be achieved by training on a single voltage imaging dataset with as few as 5000 frames. Because of the model’s self-supervised training scheme, the dataset to be denoised can also serve as the model’s only training data.

### 2.2. Architecture optimization of CellMincer via realistic physics-based simulations

To optimize the architecture and hyperparameters of CellMincer and study the impact of various design choices on the baseline denoising performance, one needs noiseless *ground truth* voltage imaging data. While experimental sourcing of true noiseless data is impractical due to technical limitations (e.g. the trade-off between signal-to-noise ratio and sampling rate, photo-bleaching and sample heating at higher illumination), we can aim to generate such ground truth recordings and their noisy realizations via carefully crafted simulations. These simulated data can then be used to study and optimize the model architecture and serve as a benchmark to evaluate the performance of CellMincer compared to other denoising methods.

To these ends, we developed Optosynth, a methodology for generating physics-based synthetic optical voltage imaging data using single-neuron morphological reconstructions and paired EP measurements from the Allen Brain Patch-seq dataset [15–17]. In brief, Optosynth simulates a *noiseless* voltage imaging readout by sampling neurons from a Patch-seq dataset, arranging them on a synthetic imaging field, and modeling the fluorescence signal density as an appropriate conversion function of the measured membrane potential. To produce realistic voltage imaging readouts, we additionally include low-passing of EP to the fluorescence sensor sampling rate, action potential wavefront propagation and decay, variability in fluorescent reporter expression, point spread function (PSF), and static and dynamics background autofluorescence. We generate noisy readings from noiseless simulations by adding Poisson shot noise and Gaussian sensor thermal noise. Optosynth’s simulations are highly customizable, enabling generation of synthetic datasets that can represent a wide range of experimental conditions, noise levels, and magnifications. A detailed description of Optosynth is provided in Supplemental Sec. S.5.

The CellMincer model is specified by a large set of hyperparameters which determine the architecture of the underlying DNNs, the self-supervised training parameters, and the optimizer scheduling. To optimize over this hyperparameter space, we first identified a baseline configuration that specifies a design empirically capable of denoising our Optosynth datasets and training it to sufficient convergence. We then constructed a series of single-hyperparameter variations of the baseline configuration and evaluated their performance on Optosynth data. Our hyperparameter variations included the inclusion or exclusion of global features *ℱ*, the length of the denoising window 2*τ* + 1, the U-Net parameters (depth, number of channels), the temporal post-processor architecture, the loss function, and the rate of pixel masking during self-supervised training. Our evaluation procedure consisted of training the model on a subset of our Optosynth data, denoising both the training data and unseen data (biological replicates generated using Optosynth) with our trained model, and computing the denoised imaging’s peak signal-to-noise ratio (PSNR) with respect to the ground truth. These results determined our final selection of hyperparameters used in subsequent benchmarking experiments.

Foremost, we found that conditioning the U-Net on global features produced the most significant improvement by wide margins, up to 5 dB gain in PSNR (see Supplementary Fig. S 2b, rows 1-3). Without inclusion of global features, model performance gains relied heavily on increasing the local denoising temporal context windows (see Supplementary Fig. S 2b, rows 1, 4-7; the context size is varied from 5 to 21 frames *∼* 10-42 ms). We note that such large temporal context sizes exceed the observed temporal correlation lengths in voltage imaging (see lagged cross-correlations in Fig. 2d and Fig. 3c), suggesting that the unconditioned denoiser is taking advantage of large context sizes to infer pixel-to-pixel spatial correlations rather than temporal correlations. To further underscore this point, we note that PSNR gains of a denoiser explicitly conditioned on precomputed global auto-correlations saturate between 5 and 9 frames *∼* 10-18 ms, which coincides with the typical temporal correlation length in neuronal activity (see Supplementary Fig. S 2e, rows 1-5). Clearly, precomputing global features and conditioning the denoiser is a much more effective and computationally efficient alternative to using longer denoising context windows.

**Figure 2.**
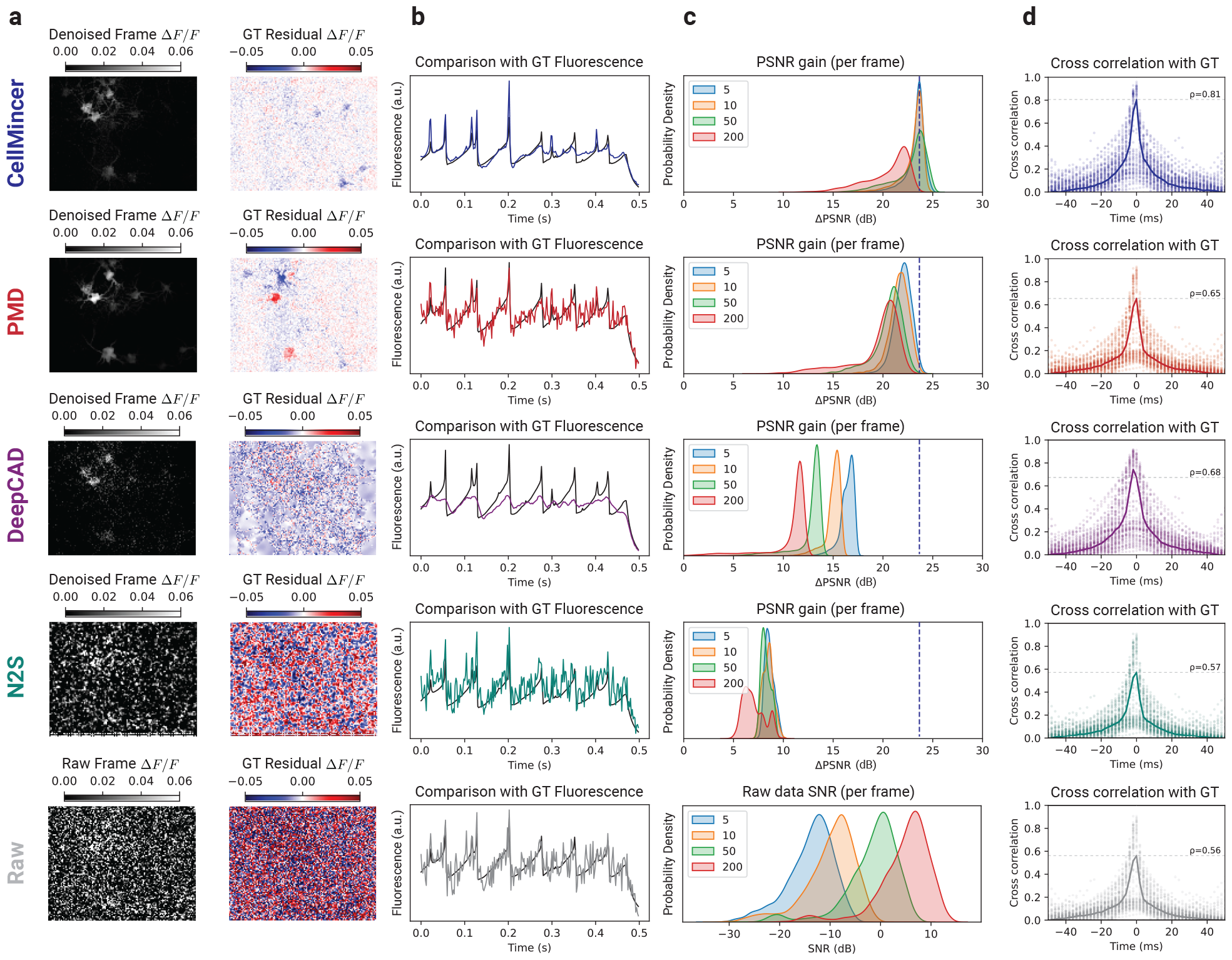
Benchmarking CellMincer and three other denoising methods on simulated voltage imaging. (a) Sample denoised frame visualizations (grayscale images) and their residuals with respect to simulated ground truth imaging (red/blue images). Both the denoised and residual images are shown as relative change in fluorescence Δ*F/F* with respect to a frame-averaged polynomial regression of the baseline (see Supplemental Sec. S.3). (b) Sample denoised ROI-averaged neuron traces (color), overlaid with the ground truth (black). (c) Distributions of single-frame PSNR gain achieved through denoising. Each distribution corresponds to a different value of simulated photon-per-fluorophore count *Q* (shown in the legend), which is the measure of raw data SNR in Optosynth simulations (see Supplemental Sec. S.5). The dashed vertical line over the top four rows is a guide for the eye and indicates the mode of CellMincer’s PSNR gain distribution for the lowest SNR data (corresponding to *Q* = 5). The plot at the bottom row shows the SNR distributions of the raw datasets at different *Q* levels. (d) Distributions of lagged cross-correlations between denoised single-neuron traces and their ground truths. Their medians are overlaid with peak correlations at Δ*t* = 0 labeled. Abbreviations: GT (ground truth).

**Figure 3.**
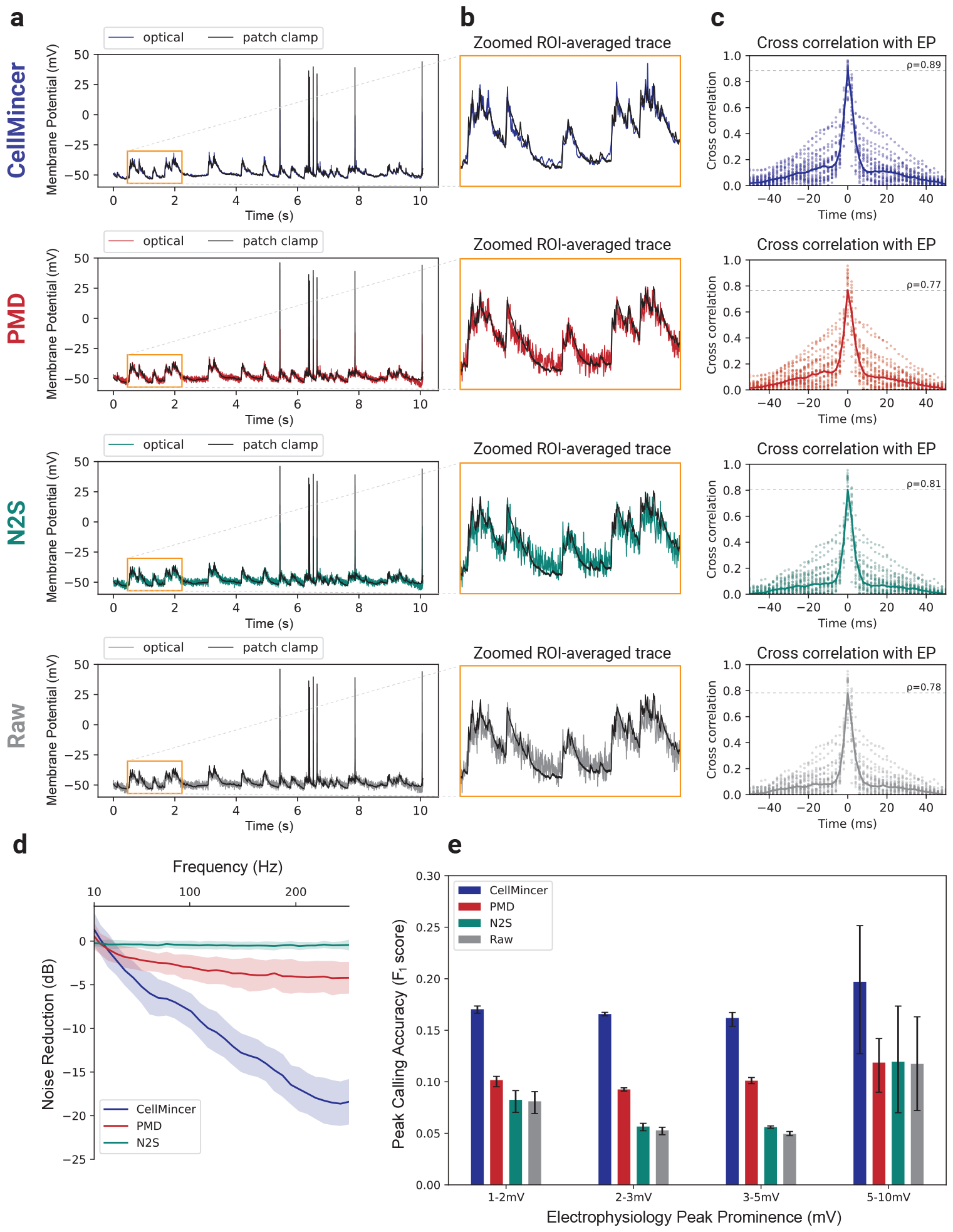
Benchmarking CellMincer and two other denoising methods on paired optical and patch clamp datasets. (a) Sample denoised ROI-averaged neuron traces (color), aligned to the EP-derived ground truth (black). (b) Inlays of subthreshold activity as indicated in the previous column, magnified. (c) Distributions of lagged cross-correlations between denoised single-neuron traces and their corresponding aligned EP signals. Their medians are overlaid with peak correlations at Δ*t* = 0 labeled. (d) Average noise reductions at varying frequency ranges achieved through denoising. (e) Peak-calling accuracy *F*_1_-scores over a range of EP peak prominence levels, using the EP signal as ground truth. Abbreviations: ROI (region of interest).

Another advantage of conditioning the denoiser on precomputed global features is achieving more robust PSNR gain characteristics across different stimulation amplitudes. This can be seen by comparing the violin plots of PSNR gain distributions for unconditioned and conditioned denoisers in Supplementary Fig. S 2. The PSNR gain distribution of the unconditioned denoiser (baseline, first row, panel b-c)) varies from +16 dB to +23 dB, and is highly variable in particular for recordings at lower electrical stimulation amplitudes (shown in blue). This observation further underscores that the performance of an unconditioned denoiser relies on its ability to infer correlations solely from the local context, which can be unreliable when neurons are not active under low stimulation. In contrast, the PSNR gain distributions of all conditioned model variants (rows 2-2, panel b-c, and all rows of panel e-f) are tightly concentrated from +22 dB to +24 dB.

Besides the crucial importance of conditioning the denoiser on global features, we find that other design decisions (like U-Net depth, number of channels, loss function selection, and the amount of masked pixels) have a surprisingly small effect on model performance. This indicates that CellMincer works reliably and does not require extensive parameter fine-tuning when used with other voltage imaging datasets. A more detailed description of these optimization experiments and their results are provided in Supplemental Sec. S.6.

### 2.3 CellMincer outperforms existing methods at denoising simulated voltage imaging data

For our benchmark evaluations, we include a selection of denoising algorithms applicable to movie-like data. Importantly, the algorithms share the precondition that clean reference data for training is not needed. DeepCAD [13] is a self-supervised deep learning algorithm for denoising calcium imaging data. Penalized matrix decomposition (PMD) [8] is a training-free algorithm based on a regularized, low-rank factorization of the data. As a baseline, we also include the original implementation of Noise2Self (N2S) [12] for images, which denoises movie frames individually.

To cover a range of noise conditions, we chose four different SNRs by varying the number of photons-per-fluorophore *Q* in simulations (see Supplemental Sec. S.5). For each level of SNR, denoted in increasing order as *Q* = 5, 10, 50, 200, we generated five synthetic datasets using Optosynth with associated ground truth. We then trained CellMincer and each of the other training-based denoising methods on three of the five datasets and subsequently used them to denoise all five datasets. PMD, which is a single-sample denoising algorithm, was used to individually denoise the five datasets instead. With these denoised datasets along with the original noisy datasets, we conducted a series of evaluations centered on comparing them to our ground truth imaging. We present our Optosynth benchmarks in Fig. 2.

To visualize imaging quality, our first benchmark evaluation compared the results of denoising a single movie frame in both absolute intensity and residual intensity with respect to ground truth, of which CellMincer and PMD exhibit notably cleaner residual frames (Fig. 2a). However, PMD retained significant error at neuron locations, while DeepCAD and N2S were less effective at removing noise throughout the frame. Our next evaluation explored the resolution of single-neuron signals from the imaging. To extract these signals, we inferred single-neuron masks from the raw data (Supplemental Sec. S.8) and used them to produce ROI-averaged traces. We overlaid these traces with their respective ground truth for a sample neuron (Fig. 2b) and observed significantly less noise from CellMincer and DeepCAD. However, we noted that DeepCAD, unlike the other methods, does not fully reconstruct the spiking events. After establishing a visual evaluation of CellMincer and the compared methods, we sought to quantify these differences with the PSNR metric. In column c, we compared the distributions of single-frame PSNR gain achieved by denoising. Each frame PSNR was computed over the union of pixels contained in the neuron ROIs gathered from all five datasets, and only frames during stimulation periods were considered. These restrictions mitigated the influence of background pixels in our performance metric. We found that CellMincer demonstrates a consistent lead in PSNR gain over the other algorithms. Additionally, CellMincer is more consistent across our range of input SNRs, while the other methods yield a clear reduction in PSNR gain for higher-quality datasets. For reference, the SNR distribution of the raw datasets are given at the bottom of column c for each of the four simulated scenarios. Finally, we aimed to show that CellMincer does indeed reconstruct the activity exhibited in the ground truth single-neuron traces without temporal bias. In column d, we computed the distributions of lagged cross-correlations between the denoised and ground truth traces over the Optosynth neurons at *Q* = 10 and overlaid the median. All cross-correlations sharply peaked at Δ*t* = 0, and CellMincer exhibited a zero-lag median cross-correlation of *ρ* = 0.81, significantly higher than of DeepCAD (0.68) and PMD (0.65).

### 2.4 CellMincer improves the detection of subthreshold events in real voltage imaging data with paired electrophysiology

To extend our evaluation of CellMincer to real data, we further evaluated CellMincer on 26 external datasets from a previously published study with simultaneous voltage imaging with chemically-synthesized voltage-sensitive fluorophore, BeRST [18, 19], and patch-clamp EP recordings, the latter of which can be repurposed as a high-confidence source of ground truth. BeRST is a chemically-synthesized far-red voltage-sensitive fluorophore. Previously, we showed that BeRST can be used in cultured rat hippocampal neurons to track changes in neuronal activity in models of development and disease [20]. Data from that study contained simultaneous recordings of BeRST fluorescence (voltage imaging) and single-cell patch clamp recordings (EP) that could serve as a ground truth for benchmarking of CellMincer. Please refer to Supplemental Sec. S.2 for the details of the experimental procedure.

Our aim in the following benchmarking experiments was to extract single-neuron denoised imaging traces and compare them to their associated patch-clamp EP signals. While both modalities operate on the same underlying neural activity, they differ substantially in sampling rate, noise characteristics, and artifacts. It is thus necessary for us to minimally resolve these data modality incompatibilities by applying a series of common filters and transformations to map them onto a shared scale. These include removing a slowly-varying trend from both measurements, temporal alignment of the two recordings, and performing a global affine transformation on the detrended fluorescence recordings to make them comparable in scale to EP recordings (in mV). See Supplemental Sec. S.8 for a detailed description of our alignment procedure.

Using an analogous approach to that presented in our simulated data benchmarking, we trained the deep learning based models on as many as all 26 of the available joint datasets, depending on the capabilities of the model implementations. DeepCAD was excluded from this benchmark due to difficulties with training it on large quantities of data, as well as its previously noted poor performance in reconstructing spiking events (see Fig. 2). With these trained models, we identified a subset of 22 datasets exhibiting discernible activity suitable for benchmarking purposes. Each model was used to denoise these benchmarking datasets, while PMD, as before, denoised each dataset individually. From the resulting denoised datasets, we extracted ROI-averaged traces and aligned them to their corresponding EP signals. The concordance between these aligned traces to the underlying EP activity forms the basis of our benchmark results, shown in Fig. 3.

Our first step was to visualize the quality of our imaging traces after mapping them to the EP scale. From plotting these traces against their corresponding EP signals (Fig. 3a), we found that CellMincer again exhibited significantly less noise than the other algorithms. The improvement is particularly apparent when examining the baseline trace relative to subthreshold activity (Fig. 3b). To characterize this noise reduction, we computed the spectral power, binned by frequency, of the residual signal before and after denoising for each algorithm (Supplemental Sec. S.9). We plot the reduction in power of these residuals, measured in dB, across the frequency range (Fig. 3d). At frequencies above 100Hz, CellMincer achieves an average noise reduction of 14.3 dB, far greater than that of PMD (3.8) and N2S (0.5). Using a process analogous to that in our simulated data benchmark, we also compared the distributions of lagged cross-correlations between the voltage imaging and EP data (Fig. 3c). CellMincer similarly exhibited a higher median cross-correlation (*ρ* = 0.89), while the other algorithms remained on par with the raw data (0.78).

After demonstrating that CellMincer, by way of its enhanced noise reduction, could potentially resolve signals of a smaller magnitude than that which can be seen by the other algorithms, we sought an approach to quantify this small-signal reconstruction fidelity. Due to difficulties with reliably processing these imaging traces with tools designed for EP signals, we devised an analytic method based on peak-calling (Supplemental Sec. S.9). From this analysis, we plotted the peak-calling accuracy of each algorithm as an *F*_1_-score, binned over several ranges of peak magnitudes (Fig. 3e), and found that CellMincer exhibited a 1.7 to 3-fold increase in *F*_1_-score over the other benchmarked algorithms and the raw data across all peak magnitude ranges. CellMincer was the only algorithm to maintain *F*_1_ score *>* 0.15 across all ground truth mV changes (from 1 mV to 10 mV). At the 5-10 mV range, in contrast to CellMincer, the other two denoising methods (PMD, N2S) do not show any statistically significant improvement over the raw data. At changes below 5 mV, only CellMincer and PMD improve the accuracy over the raw data while N2S does not. Furthermore, CellMincer outperforms the accuracy of PMD significantly in this regime. These evaluations demonstrated that on real voltage imaging, CellMincer produces significant quantitative improvements in the reconstruction of single-neuron traces when compared to traces derived from the raw data and from data denoised by standard algorithms. These improvements have meaningful implications for the potential uses of CellMincer denoising to recover underlying subthreshold activity using voltage imaging data.

### 2.5 CellMincer improves neuron segmentation and detection of subtle changes in neural activity

To demonstrate the utility of CellMincer in a representative end-to-end biological hypothesis testing workflow, we present a complete such analysis with and without CellMincer as a data-denoising component, and we quantify the impact of CellMincer on improving the detection of subtle phenotypic changes. Specifically, we compare the spiking activity of unperturbed and chronically tetrodotoxin (TTX)-treated cultured hPSC-derived neurons via Optopatch voltage imaging. TTX is a voltage-gated sodium channel blocker which, when used to treat cultured neurons, prevents them from firing action potentials. Prolonged silencing with TTX increases intrinsic excitability of neurons [21]. This homeostatic plasticity is also displayed in hPSC-derived neurons [3]. We incubated hPSC-derived neuronal cultures in 500 nM TTX for 48 hours and washed it out prior to Optopatch recordings. Parallel unperturbed cultures were incubated in TTX-free media. In both cases, we subjected the neurons to eight stimulation periods in increasing intensity and measured their action potential via Optopatch voltage imaging. Please refer to Supplemental Sec. S.1 for the details of the experimental procedure.

We analysed the obtained recordings as follows. We performed a pixelwise detrending preprocessing step on both raw and denoised datasets, computed independent spatial components using a PCA/ICA decomposition approach [22], and identified neuronal components through careful comparative review of the obtained components and the activity traces. We finally derived an ROI-averaged trace from each identified neuron for downstream analysis, which focused on counting and comparing the statistics of high-amplitude action potential spikes. Additional details are provided in Supplemental Sec. S.10.

Fig. 4a showcases a segmentation of identifiable neurons on a sample frame from the raw and CellMincer-denoised analysis. It is evident that: (1) among the neurons that are reliably identifiable in both the raw and denoised dataset, CellMincer more clearly delineates their boundaries. This is particularly evident among the cluster of overlapping neurons shown on the left side of the field of view in Fig. 4a); (2) CellMincer enables better separation and detection of more neuronal components, in particular neurons with fainter fluorescence signal, as well as more reliable spike-counting. To substantiate the latter, we plotted the ROI-averaged traces from three neurons side-by-side in Fig. 4b (color-matched to the neuron components shown in Fig. 4a). As explored in the previous experiments, the most salient improvement brought about by CellMincer is in the form of a significant reduction in the background noise (see Fig. 3c). In the present context, this has the effect of highlighting subtler spiking events and enabling them to be called with greater confidence (compare the raw and denoised traces shown in Fig. 4b). The total number of confidently detected neurons are shown in Fig. 4d and establishes that in most recordings, CellMincer denoising allows identification of twice or more as many neurons with distinct spiking patterns. As a result of improved neuron segmentation and spike counting following denoising, aggregating spike statistics over more neurons results in a larger statistical separation between the control and chronically TTX-treated populations. This can be visualized by comparing the boxplots shown in Fig. 4c. To further quantify this finding, we performed a Wilcoxon rank sum test, the result of which is shown in Fig. 4e. Notably, CellMincer denoising yields significantly greater statistical power to separate the two conditions, with this separation increasing at higher stimulation intensities, as evidenced in the tail-end of Fig. 4e. Interestingly, the lowest stimulation intensity shows a deviation from this trend, yet it still aligns with the overall conclusion that chronically TTX-treated neurons exhibit heightened excitability.

**Figure 4.**
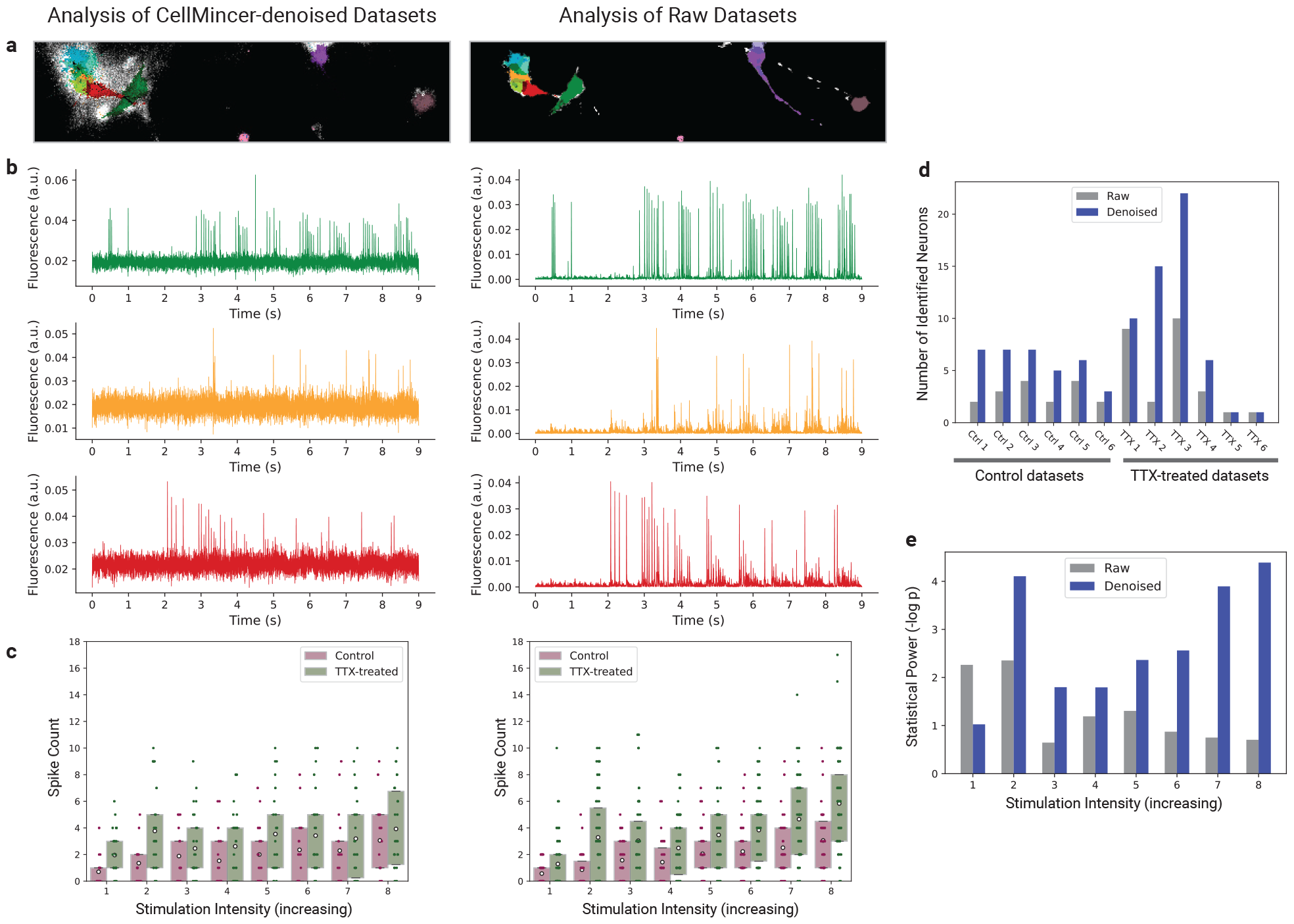
Comparing the spiking activity of chronically tetrodotoxin (TTX)-treated vs. control hPSC-derived neurons with raw and CellMincer-denoised Optopatch voltage imaging data. (a) Raw and denoised versions of a sample frame, colored with the neuron components identified in their corresponding datasets. (b) Corresponding ROI-averaged single-neuron traces detected in both versions of the above frame. (c) Spike count distributions, separated by neuron population and stimulation intensity. Spikes were identified in each detected neuron’s trace and binned by their stimulation intensity. (d) Detected neuron counts in the raw and denoised versions of each dataset. (e) Statistical power of the Wilcoxon Rank Sum test applied to the neuron population differentiation hypothesis, reported as the negative logarithm of its p-value.

## 3 Discussion

We introduced CellMincer, a self-supervised deep learning method specifically designed for denoising voltage imaging datasets, and discussed several key methodological refinements over the existing approaches. These include: (1) an efficient and expressive two-stage spatiotemporal data processing deep neural network architecture, comprising a frame-wise 2D U-Net module for spatial feature extraction, followed by a pixelwise 1D convolutional module for temporal data post-processing; (2) performing self-supervised training by masking a sparse set of pixels rather than entire frames, allowing the model to access both intra- and inter-frame information as needed for effective denoising of voltage imaging datasets; (3) conditioning local denoisers on a set of precomputed spatiotemporal auto-correlations at multiple length scales, resulting in a significant boost in denoising accuracy; (4) introducing a physics-based simulation framework to generate highly realistic pairs of clean and noisy voltage imaging movies for the purpose of hyperparameter optimization and ablation studies. We evaluated CellMincer’s performance on both simulated and real datasets, including an external previously published dataset comprising 30 voltage imaging experiments with simultaneous patch-clamp EP recordings [20], and established that CellMincer outperforms the existing denoising approaches. Finally, we demonstrated the utility of CellMincer in downstream analyses, resulting in a more robust identification of neurons and spiking events and ultimately a higher statistical power for separating neuron populations based on functional phenotypes.

CellMincer denoising holds the potential to advance the study of complex neuronal communication through multiple avenues. Firstly, traditional methods often involve measuring postsynaptic potentials at the cell body to understand synaptic transmission. However, the inherent biophysical properties of synapses, coupled with the intricate dendritic morphology, such as shape, branching, and diameter, can distort electrical signals originating at the synapse. Consequently, the activity recorded at the soma may not accurately depict events at the synapse. This discrepancy poses a challenge in electrophysiological techniques, which predominantly require recordings at the soma. CellMincer, however, presents a promising solution by facilitating the direct examination of electrical activity at the synapse through voltage imaging techniques and computational SNR enhancement. Secondly, CellMincer denoising enables the collection of usable data even from low magnification recordings with low signal-to-noise ratios (SNRs). For instance, enough data was acquired in seconds to clearly separate the TTX-treated groups (Fig. 4). This is in contrast to single cell recordings either by patch-clamp EP or high magnification voltage imaging, which would take multiple recording days.

CellMincer’s improved performance over similar denoising methods largely stems from precomputing global movie features and using these features as a *conditioner* for the denoiser network that otherwise operates on small temporal contexts. The inclusion of global spatial features alongside CellMincer’s localized context processing allows the algorithm to exploit persistent long-range correlations in the data without directly ingesting the entire dataset with a neural network, an intractable computational operation. These carefully crafted features (see Supplemental Sec. S.3) meaningfully contribute to data modeling because the neuron signal sources are generally fixed in space, making the behavior of individual pixels, as well as pixel-pixel relationships, highly consistent. Our ablation studies reveal that providing global features to the denoiser results in a striking 3-fold boost in the PSNR (or approximately a 5 dB gain). While increasing the denoising context size endows the local denoiser with more global information and improves its performance, we notice that there is still a *∼* 4 dB gap between an unconditioned denoiser with a context size of 21 frames and a conditioned denoiser with a context size of only 9 frames, see Supplementary Fig. S 2.

Another advantage of leveraging precomputing global features in conditioning short-context denoisers is computational efficiency. Existing denoising methods that rely on long contexts to achieve satisfactory denoising performance will inevitably demand relatively larger computational resources. For instance, when extended to a training corpora of 26 datasets (Sec. 2.4), DeepCAD’s computational resource demands exceeded our limits and we had to exclude it from the benchmarking. Likewise, DeepInterpolation [14], another self-supervised deep neural network denoiser, could not be made to process our datasets and was thus also not included in benchmarking. In contrast, we were able to efficiently train CellMincer on our largest training corpora using widely-available commodity GPUs.

The pre-training approach we employed to train the CellMincer model on voltage imaging data might not be optimally tailored to handle other functional imaging modalities, such as calcium imaging, characterized by substantially different spatiotemporal dynamics. A characteristic difference in the dynamics of voltage imaging and calcium imaging is the presence of single spiking events occurring within 5 to 10 frames. The performance gap between CellMincer and DeepCAD on voltage imaging suggests that the most effective fluorescence imaging denoisers are highly specific to their target domains. CellMincer’s architecture uses a context window length on par with the timescale of a typical voltage imaging spiking event, and its training scheme maximizes the utility of this context by enabling inference from same-frame pixels. Conversely, DeepCAD predicts whole frames from a large, temporally downsampled neighborhood of frames, a strategy which foregoes mutual information carried by proximal pixels in exchange for training scalability. Our hypothesis of the specificity of the model (and self-supervision task) to the data domain is further reinforced by an experiment in which we compare the performance of CellMincer and DeepCAD on calcium imaging data (see Supplementary Fig. S 5). Both algorithms were trained on seven low-SNR calcium imaging datasets [13], and their denoised outputs were compared to high-SNR versions of the same datasets. We noticed a significant drop in the performance of CellMincer on denoising calcium imaging data, and improved performance of DeepCAD, which is opposite to our findings on fast voltage imaging data. We concluded that CellMincer’s capacity to model short-term fluctuations becomes a hindrance when the underlying signal has inherently slow dynamics, whereas DeepCAD’s whole-frame masking and the implicit bias of slow dynamics becomes advantageous.

A notable difficulty in conducting the analysis of neuron imaging traces in relation to corresponding EP activity is the lack of available tools for analyzing waveforms that diverge from the highly specific characteristics of EP. The Electrophys Feature Extraction Library (EFEL) [23], one such tool, can extract a variety of EP features such as spike half-widths but is much less conclusive when the input signals were adapted from fluorescence imaging. Our solution, prominence-thresholded peak calling, is motivated by the biological significance of partial and total depolarization events, indicative of subthreshold activity and action potentials respectively. Thus, identifying peaks in the EP represents a sensible first-order approximation for the locations of these events and can function as a task to which we subject our imaging traces. While most action potentials stand in such stark contrast to the surrounding baseline that they are evident in any form of the trace, the elevated baseline in the traces produced by PMD, N2S, and the raw data is likely to hide less pronounced EP events and introduce more false positives. Although the absolute peak-calling performance across all methods is low, primarily owing to the inherent incompatibilities between the 50 kHz EP signal and its derived 500 Hz imaging, our assessment is that for EP peaks between 2-5 mV, there is indeed information in the raw data that corresponds to this activity but is not immediately visible, and CellMincer is distilling this information to allow for more confident judgements.

A limitation of CellMincer’s default self-supervised training scheme is that in uniformly sampling random crops of the training data, CellMincer spends an overwhelming majority of its computation time on static background pixels as opposed to pixels containing meaningful neuronal activity. We introduced an option in CellMincer to increase its sampling efficiency without introducing network bias by oversampling such *meaningful* data crops, defined as exceeding the top *n*% of average luminosity across all crops in the dataset for *n* chosen between 0.1 and 1, to 50% of each training batch (importance sampling). We can then correct the loss calculation knowing the constructed ratio of meaningful samples. While this feature was not incorporated into the models used in our main benchmarking experiments, we found that it reduced the performance gap between CellMincer and DeepCAD on calcium imaging. We expect further exploration of this direction, namely adaptive sampling and *hard sample* mining in the context of self-supervised training, will find application beyond the present domain.

Precomputation of global features should be carried out with additional considerations in situations where either neuron locations or noise characteristics could be non-stationary (e.g. certain *in vivo* recordings). In such cases, the temporal distributional shift along a long recording interval may render any one set of precomputed *global* features less relevant to variable local contexts of the recording. Since CellMincer’s objective function is unbiased (see Eq. 4), conditioning on poor (or even irrelevant) global features will not degrade the performance of the method. However, to take full advantage of the notion of feature-conditioning, we stipulate that a more effective strategy in denoising non-stationary data would be to pre-segment the movie into approximately stationary intervals and denoising each section separately using its own precomputed features. This is streamlined by CellMincer’s ability to pre-train on and denoise an arbitrary collection of recordings.

We believe that CellMincer’s architecture is adequately powered for denoising many forms of fluorescence imaging modalities of electrically active cells. Optimizing the hyperparameters and training schedule of CellMincer for related data domains (e.g. calcium imaging) would be a natural avenue for future work. While our analysis shows that CellMincer satisfactorily operates on a single dataset (both for training and denoising), we hypothesize that training a generalist large-scale CellMincer *foundation model* on a large and diverse biomedical imaging corpora (and perhaps using more scalable architectures such as the Vision Transformer [24]) is another promising area of future research. Intriguingly, inspecting and building on the saliency and attention maps underlying the Vision Transformer could lay a novel roadmap for segmenting a wide range of functional imaging datasets into functional units, much like the recently demonstrated utility of self-supervised models of natural images (e.g. DINOv2 [25]) in segmenting natural images and performing various other image-based downstream tasks.

We have made available separate pre-trained CellMincer models on synthetic Optosynth data from Sec. 2.3, the BeRST voltage imaging data from Sec. 2.4, and the Optopatch voltage imaging data from Sec. 2.5. Even though training a CellMincer model from scratch can take 10-12 hours on a typical dataset and publicly available commodity GPU (see Supplemental Sec. S.4), using one of the pre-trained models as is or fine-tuning it presents a faster and less computationally intensive approach to the adoption of our method. In the future, we hope that the availability pretrained CellMincer foundation models on a large and diverse voltage imaging dataset, combined with efficient model selection and fine-tuning strategies, will further reduce the computational cost of using CellMincer.

## 4 Acknowledgements

This work was supported by a grant from SFARI (#890477, SLF, RN), by the National Institute of Mental Health (R01 MH128366-01A1, SLF, RN; R21 MH120423, SLF; and U01MH115727, MB, RN), by the National Institute of Health (R01 NS098088, ASW, EWM), by the Stanley Center for Psychiatric Research, by a SPARC award from Broad Institute (year 2019-2020, TM, MB, SLF), and by a BroadIgnite award from Broad Institute (2021-2022, BW, MB). The template HT076 was a gift from Dr. Adam Cohen. The authors thank Luca D’Alessio for insightful comments during various stages of the development and testing of CellMincer.

## 5 Code Availability

The code repository containing the CellMincer pipeline can be found at https://github.com/cellarium-ai/CellMincer. The repository containing the Optosynth simulation framework can be found at https://github.com/cellarium-ai/Optosynth. An auxiliary repository with reproductions of the analysis used to produce the figures can be found at https://github.com/cellarium-ai/CellMincerPaperAnalysis.

## 6 Data Availability

All data used to conduct the benchmarking experiments with Optosynth data, BeRST voltage imaging with paired EP data, and Optopatch datasets for chronically TTX-treated and unperturbed hPSC-derived neurons can be accessed either directly from the Google Cloud bucket found at gs://broad-dsp-cellmincer-data, or through notebooks in the aforementioned paper analysis repository.

## S Supplemental Sections

### S.1 Optopatch voltage imaging of chronically TTX-treated and unperturbed hPSC-derived neurons

hPSC-derived neurons differentiation was performed as previously described [26, 27], followed by Optopatch voltage imaging also as previously described [28]. Methods are reproduced below for completeness.

#### hPSC Culture

Human ESCs were maintained on plates coated with Geltrex (Life Technologies, A1413301) in mTeSR Plus medium (StemCell Technologies, 100-1130) and passaged with Accutase (Gibco, A11105). All cell cultures were maintained at 37 ^*°*^C, 5% CO2.

#### Neuronal Induction

hPSC-derived neurons were differentiated from an hPSC line (H1/WA01) using combined NGN2 programming with SMAD and WNT inhibition in the presence of mouse astrocytes [26, 27]. On day 0, hPSCs were differentiated in N2 medium (500 mL DMEM/F12 [1:1] [Gibco, 11320-033]), 5 mL Glutamax (Gibco, 35050-061), 7.5 mL sucrose (20%, Sigma, S0389), 5 mL N2 supplement B (StemCell Technologies, 07156) supplemented with SB431542 (10 μM, Tocris, 1614), XAV939 (2 μM, Stemgent, 04-00046), and LDN-193189 (100 nM, Stemgent, 04-0074) along with doxycycline hyclate (2 μg.mL^*−*1^, Sigma, D9891) and Y27632 (5 mM, Stemgent 04-0012). On day 1 and 2 media was changed to N2 medium supplemented with SB431542 (5 μM, Tocris, 1614), XAV939 (1 μM, Stemgent, 04-00046), and LDN-193189 (50 nM, Stemgent, 04-0074) with doxycycline hyclate (2 μg.mL^*−*1^, Sigma, D9891) and Zeocin (1 μg.mL^−1^, Invitrogen, 46-059). On Day 3 neuronal precursor cells were passaged with Accutase into Neurobasal media (500 mL Neurobasal [Gibco, 21103-049], 5 mL Glutamax [Gibco, 35050-061], 7.5 mL Sucrose [20%, Sigma, S0389], 2.5 mL NEAA [Corning, 25-0250 Cl]) supplemented with B27 (50x, Gibco, 17504-044), BDNF, CTNF, GDNF (10 ng.mL^*−*1^, R&D Systems 248-BD/CF, 257-NT/CF, and 212-GD/CF) and doxycycline hyclate (2 μg.mL^*−*1^, Sigma, D9891) in a 24-well format and infected with lentiviral optogenetic constructs (HT076, hSyn Cre-off Archon-TS-darkCitrine-TSx3-ER) at 2 MOI. On day 7, the cells were passaged with Accutase onto 10mm glass coverslip bottom dishes precoated with Geltrex containing a monolayer of mouse cortical astrocytes. Estimated neuron/astrocyte ratio was 1:2 with 80k neurons plated per 10mm dish. Cell were matured in Neurobasal media supplemented with B27 (50x, Gibco, 17504-044), BDNF, CTNF, GDNF (10 ng.mL^*−*1^, R&D Systems 248-BD/CF, 257-NT/CF, and 212-GD/CF) and doxycycline hyclate (2 μg.mL^−1^, Sigma, D9891) with 50% media changes twice a week.

#### TTX treatment and Optopatch imaging

On Day 35 500 nM TTX (Tocris) was added to the culture media and cultures were returned to the incubator for additional 48 hours. Parallel control cultures were kept in TTX-free media. 10 minutes prior to recording the cultures were washed 3 times in pre-warned recording solution (125 mM NaCl, 2.5 mM KCl, 3 mM CaCl2, 1 mM MgCl2, 15 mM HEPES, 30 mM glucose (pH 7.3) and adjusted to 305–310 mOsm with sucrose) to wash out the TTX. Recordings were obtained in recording solution in the absence of TTX at 23 ^*°*^C. Cellular activity was recorded on custom-built wide-field microscope equipped with oblique illumination lens and a wide 20x objective. The cells were stimulated with 500 ms blue light (488 nm) at 1 Hz of increasing intensity (20 to 120 mW/cm^2^) for 6 seconds, while firing patterns were recorded under continuous red light (635 nm) illumination at 1 kHz.

### S.2 Simultaneous BeRST fluorescence voltage imaging and single-cell patch-clamp EP recording experimental procedure

Simultaneous BeRST imaging and single-cell patch-clamp EP recordings were performed as described previously [20]. Methods are reproduced below for completeness.

#### Cell Culture

All animal procedures were approved by the UC Berkeley Animal Care and Use Committees and conformed to the NIH Guide for the Care and Use of Laboratory Animals and the Public Health Policy.

#### Rat Hippocampal Neurons

Hippocampi were dissected from embryonic day 18 Sprague Dawley rats (Charles River Laboratory) in cold sterile HBSS (zero Ca2+, zero Mg2+). All dissection products were supplied by Invitrogen, unless otherwise stated. Hippocampal tissue was treated with trypsin (2.5%) for 15 min at 37 ^*°*^C. The tissue was triturated using fire polished Pasteur pipettes, in minimum essential media (MEM) supplemented with 5% fetal bovine serum (FBS; Thermo Scientific), 2% B-27, 2% 1 M D-glucose (Fisher Scientific) and 1% GlutaMax. The dissociated cells (neurons and glia) were plated onto 12 mm diameter coverslips (Fisher Scientific) pre-treated with PDL at a density of 30-40,000 cells per coverslip in MEM supplemented media (as above). Cells were maintained at 37 ^*°*^C in a humidified incubator with 5% CO2. At 1 day in vitro (DIV), half of the MEM supplemented media was removed and replaced with FBS-free media to supress glial cell growth (Neurobasal media containing 2% B-27 supplement and 1% GlutaMax). Functional imaging was performed on 8-15 DIV neurons to access neuronal excitability and connectivity across different stages of development. References to biological replicates, or “n,” refer to the number of dissections data were collected from.

#### VoltageFluor/BeRST 1 Stocks and Cellular Loading

For all imaging experiments, BeRST 1 was diluted from a 250 μM DMSO stock solution to 0.1-1μM in HBSS (+Ca2+, +Mg2+, -phenol red). To load cells with dye solution, the media was first removed from a coverslip and then replaced with the BeRST-HBSS solution. The dye was then allowed to load onto the cells for 20 minutes at 37 ^*°*^C in a humidified incubator with 5% CO2. After dye loading, coverslips were removed from the incubator and placed into an Attofluor cell chamber filled with fresh HBSS for functional imaging at room temperature (20-23 ^*°*^C).

#### Voltage Imaging with BeRST

Voltage imaging was performed on an upright AxioExaminer Z-1 (Zeiss) or an inverted Zeiss AxioObserver Z-1 (Zeiss), both equipped with a Spectra-X light engine LED light (Lumencor), and controlled with Slidebook (3i). Images were acquired using a W-Plan-Apo/1.0 NA 20x water immersion objective (Zeiss) or a Plan-Apochromat/0.8 NA 20x air objective (Zeiss). Images (2048 px *×* 400 px, pixel size: 0.325 μm *×* 0.325 μm) were collected continuously on an OrcaFlash4.0 sCMOS camera (sCMOS; Hamamatsu) at a sampling rate of 0.5 kHz, with 4*×*4 binning, and a 631 nm LED (13 mW/mm^2^, SpectraX) with a 631/28 nm excitation bandpass. Emission was collected after passing through a quadruple bandpass dichroic (432/38 nm, 509/22 nm, 586/40 nm, 654 nm LP and quadruple bandpass emission filter (430/32 nm, 508/14 nm, 586/30 nm, 708/98 nm).

#### Electrophysiology

For electrophysiological experiments, pipettes were pulled from borosilicate glass (Sutter Instruments, BF150-86-10), with a resistance of 5–8 MΩ, and were filled with an internal solution; (in mM) 115 potassium gluconate, 10 BAPTA tetrapotassium salt, 10 HEPES, 5 NaCl, 10 KCl, 2 ATP disodium salt, 0.3 GTP trisodium salt (pH 7.25, 275 mOsm). Recordings were obtained with an Axopatch 200B amplifier (Molecular Devices) at room temperature. The signals were digitized with a Digidata 1440A, sampled at 50 kHz and recorded with pCLAMP 10 software (Molecular Devices) on a PC. Fast capacitance was compensated in the on-cell configuration. For all electrophysiology experiments, recordings were only pursued if the series resistance in voltage clamp was less than 30 MΩ. For whole-cell, current clamp recordings in hippocampal neurons, following membrane rupture, resting membrane potential was assessed and recorded at *I* = 0 and monitored during the data acquisition.

### S.3 CellMincer preprocessing and global feature extraction details

Before a voltage imaging movie *X*(*t, x, y*) is received as input to a CellMincer model, our pipeline applies several preprocessing steps to: (1) approximately isolate the background fluorescence; (2) normalize the dynamic range of background-subtracted data prior to denoising; (3) precompute a number of global movie statistics for conditioning the local denoiser. In this section, we detail the data preprocessing and global feature extraction stages of the CellMincer pipeline.

#### Data preprocessing and trend isolation

Background fluorescence is a dynamic imaging artifact both highly individual to its source dataset and magnitudes larger than the true fluorescence signal, so removing it aids the network in identifying neuron action potentials. To model this background activity separately for each (*x, y*) pixel, we temporally interpolate each pixel’s trace with a low-order polynomial (with a default value of *n*_poly_ = 3) to obtain the following decomposition:

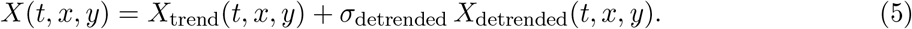

By design, *X*_trend_ approximately captures the smooth temporal trend and DC bias offset in the recording, whereas *X*_detrended_ represents the normalized residual fluorescence signal. When specified by the user, we obtain the smooth trend fit only from the resting periods (typically the beginning and the end segments of a recording segment). When such resting periods are not included in the recording, we regress over the entire recording and use a lower order polynomial (*n*_poly_ = 1) to avoid overfitting to the neural activity. We note that normalizing the detrended component by its standard deviation over all pixels and time points, *σ*_detrended_, allows CellMincer to train over multiple datasets and data sources (see Eq. 5). After denoising such a detrended dataset, CellMincer reports both the output without modification and reconstituted with the original scaling and trend (Eq. 5).

#### Precomputing global features

After the preprocessing stage of the CellMincer pipeline, we precompute the global features as follows. First, we further decompose *X*_detrended_(*t, x, y*) into slow and fast components:

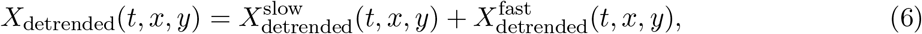

where 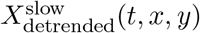 is the moving average of *X*_detrended_(*t, x, y*) over a short window. For a 500 Hz recording, we calculate the moving average over 10 frames, corresponding to 20 ms. The goal here is to separate the neural activity into fast transients (e.g. spikes) and slower features (e.g. subthreshold activity). We calculate the same set of global features from the two components independently. We define the general spatially-resolved temporal auto-correlation function as such:

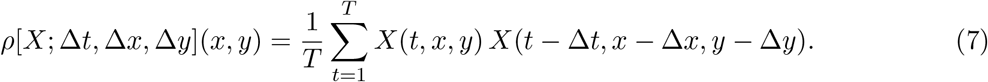

The first three global features are: (1) the square root of 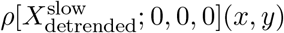, i.e. the pixelwise slow temporal variability; (2) the square root of 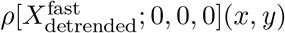, i.e. the pixelwise fast temporal variability; (3) the temporal mean of 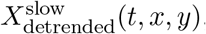, i.e. the mean pixelwise slow activity. In addition to these, we include 17 other normalized and spatially-resolved auto-correlation functions as follows:

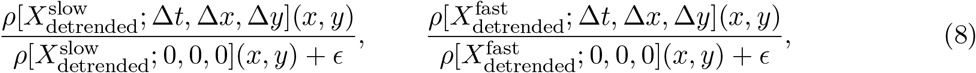

for Δ*x*, Δ*y* ∈ {−1, 0, 1}, Δ*t* ∈ {0, 1}, and excluding Δ*x* = Δ*y* = Δ*t* = 0. Put together, these amount to 37 feature maps. Next, we spatially downsample both 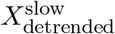 and *X* 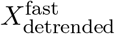 by a factor of two, such that each image-space pixel corresponds to the average signal over a two native-resolution pixels. We calculate the same set of 37 features maps, and upsample the obtained feature map by a factor of two back to the original resolution. The rationale is to bring more distant spatially-lagged auto-correlations into a feature map in the native resolution. In principle, this procedure can be repeated multiple times to capture further dilated and averaged auto-correlations. We stop the procedure at the second level, obtaining *F* = 2 *×* 37 = 74 spatial feature maps in total which we collect and concatenate into a *F × H × W* tensor. Conveniently, the *F* channels of this tensor encode a standardized set of spatiotemporal auto-correlations at different lengthscales, which can be used by the model to infer covarying groups of pixels without having access to the full movie.

### S.4 CellMincer neural network design, training schedule, and implementation details

The neural network architecture of CellMincer consists of a U-Net which produces deep embeddings of individual frames and a temporal post-processor which reduces a sequence of frame embeddings into a single denoised frame (see Fig. 1b).

Our U-Net design allows for the augmentation of the input frame with our precomputed global features. This augmentation can occur either by concatenating the two before passing it through the U-Net or by repeating this concatenation at each step of the U-Net’s contracting path, iteratively downsampling the global features in tandem with the frame embedding (see Fig. 1). We find that this *repeat* global feature augmentation reinforces the network bias toward using the global features, improving downstream performance. In addition, our U-Net implementation is not limited to a specific input dimension (as often required by conventional implementations), as demonstrated by our protocol of training on small imaging crops while using whole frames at evaluation time. This allows the model to train on imaging corpora with mixed dimensions and generalize to arbitrarily sized inputs without needing to dissect the input into uniformly-sized patches. Without padding each convolution layer, our U-Net produces image contraction, so we apply reflection padding to the input to achieve our desired output dimensions.

The temporal post-processor takes as input a short window of frame embeddings from the U-Net, convolves the time dimension, and collapses the feature dimension, producing a single output for each pixel. In this manner, no further spatial entanglement is introduced, so we do not include global features at this computation step (see Fig. 1d).

Through optimization trials, we determined an Adam optimizer with standard momentum parameters (*β*_1_ = 0.9, *β*_2_ = 0.999) was most effective for training CellMincer. We applied a cosine-annealed learning rate with linear warmup [29], parameterized at *η*_max_ = 10^*−*4^. To increase the diversity of imaging used to train our model, we configured our training samples to consist of small 62*×*62 crops padded with 30 pixels on each side, striking a balance between the minibatch diversity and the training signal that comes from each entry in the minibatch. With this configuration, we were able to maximize GPU utilization by training on minibatches of 20 samples per GPU (reduced to 10 samples for our largest model variant). We found that 50,000 training iterations generally led to sufficient model convergence when using a training set of limited size (1-5 recordings). More investigation is needed to determine whether a longer training period is needed to make full use of a larger training set.

In the course of CellMincer’s development, we explored a series of variations on its architecture and training schemes, some of which are reported in detail in Sec. S.6. Of those omitted from our results, we considered single U-Net architectures that combined spatial and temporal processing, either by using a 3D U-Net to model time or by concatenating the frame sequence within the feature dimension. While our *time as features* model was computationally more efficient, we could not reach the expressivity and performance afforded by our current two-stage design. We also experimented with the choice of learning rate schedule, opting for an empirically validated cosine annealing with 10% linear warmup [29]. This schedule mitigates early instability while the model performance is highly variant while improving convergence near the end of training. In addition, we briefly explored the use of stochastic weight averaging [30], and discovered that it had a paradoxically negative impact on performance. Our hypothesis is that the contours of our model’s loss landscape are highly nonconvex so that an averaging of local optima removes us from the optimal parameter manifold.

We implemented CellMincer as a CLI tool in the PyTorch Lightning framework, which offers ease of scalability with multi-GPU training. To offer a sense of training costs and runtime, a CellMincer model trained on a single dataset using one NVIDIA Tesla T4 GPU for a standard 50,000 iterations would take 12-16 hours to finish, while a larger operation using 26 datasets and 4 GPUs may take 6-7 days. In practice, we found that training a fresh model to denoise a new dataset is not necessary. For instance, in our end-to-end hypothesis testing experiment (Sec. S.10), we simply used a model previously trained on a different voltage imaging corpus (recorded under similar conditions) and did not involve any of the datasets in the experiment. Denoising a typical dataset with CellMincer, by contrast, takes no more than a few minutes on any GPU setup.

### S.5 Simulating realistic voltage imaging datasets using Optosynth

In order to optimize the architecture and hyperparameters of a data denoising technique and study the impact of various design choices on the bottom line denoising performance, one needs noiseless or high-SNR *ground truth* data. To generate such ground truth recordings and their noisy realizations, we developed a physics-based simulation companion software called *Optosynth* in which we aimed to carefully model salient aspects of the phenomenology of voltage imaging. We briefly describe the key steps involved in Optosynth simulations as follows (see Supplementary Fig. S 1 for a graphical overview).

#### Data procurement and preprocessing

We used Allen SDK [31] to access Allen Brain Atlas data, and procured 485 neurons from mouse primary visual cortex (VISp) from the Allen cell types database with paired morphology and EP data (Patch-seq) [32–34]. We minimally preprocess reconstructed morphology data as follows. We scale the image-space pixel from the original 0.1144 μm/pixel by a factor of 10 to 1.144 μm/pixel, representative of the typical magnification of voltage imaging experiments. We project the 3D morphology into a 2D binary mask. Allen morphology reconstructions only provide the location and radius of the soma. We use this information to generate a synthetic soma shape circumscribed by a random Fourier curve with 3 frequency components and a wiggle amplitude of no more than 20% of the soma radius. We also minimally preprocess the EP data by truncating the sweeps to 2 sec in duration, amounting to 0.5 sec of resting recording, 1s of electrical stimulation, followed by another 0.5 sec of resting recording. For each neuron, we keep track of the stimulation current amplitude (pA) and the membrane potential (mV) time series.

#### Setting up the simulation

The Optosynth simulation configuration is used to generate a manifest for the experiment. The key user input is the number of desired stimulation segments for the full experiment, and is given to the tool as a list of [*I*_min_, *I*_max_] stimulation amplitude intervals. Based on this specification, we filter the pool of available Patch-seq neurons to the subset of neurons that have at least one sweep within each stimulation interval. Relaxing the stimulation current to fall within an interval (in contrast with matching a specific value) allows more flexibility in choosing neurons for simulation, because not all Patch-seq neurons have received the same set of stimulation amplitudes.

#### Simulating the neuron fluorescence density

For each neuron, we precompute a spike decay map *α*(**r**), a delay map *τ* (**r**), and a fluorescent reporter spatial density map *ρ*(**r**) as follows:

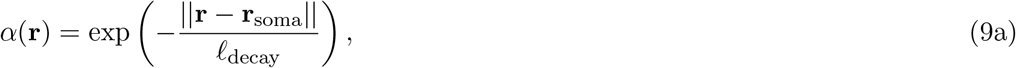

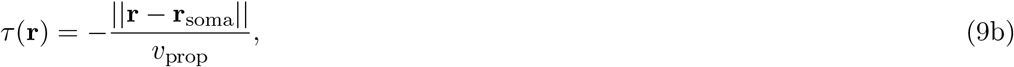

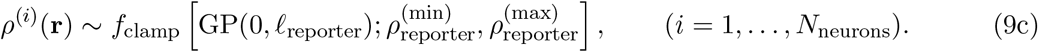

Here, **r** is a 2D position vector, **r**_soma_ refers to the soma location of the neuron of interest, *𝓁*_decay_ is a specified signal decay lengthscale, *v*_prop_ is the action potential propagation velocity [3, 35].

**Figure S 1:**
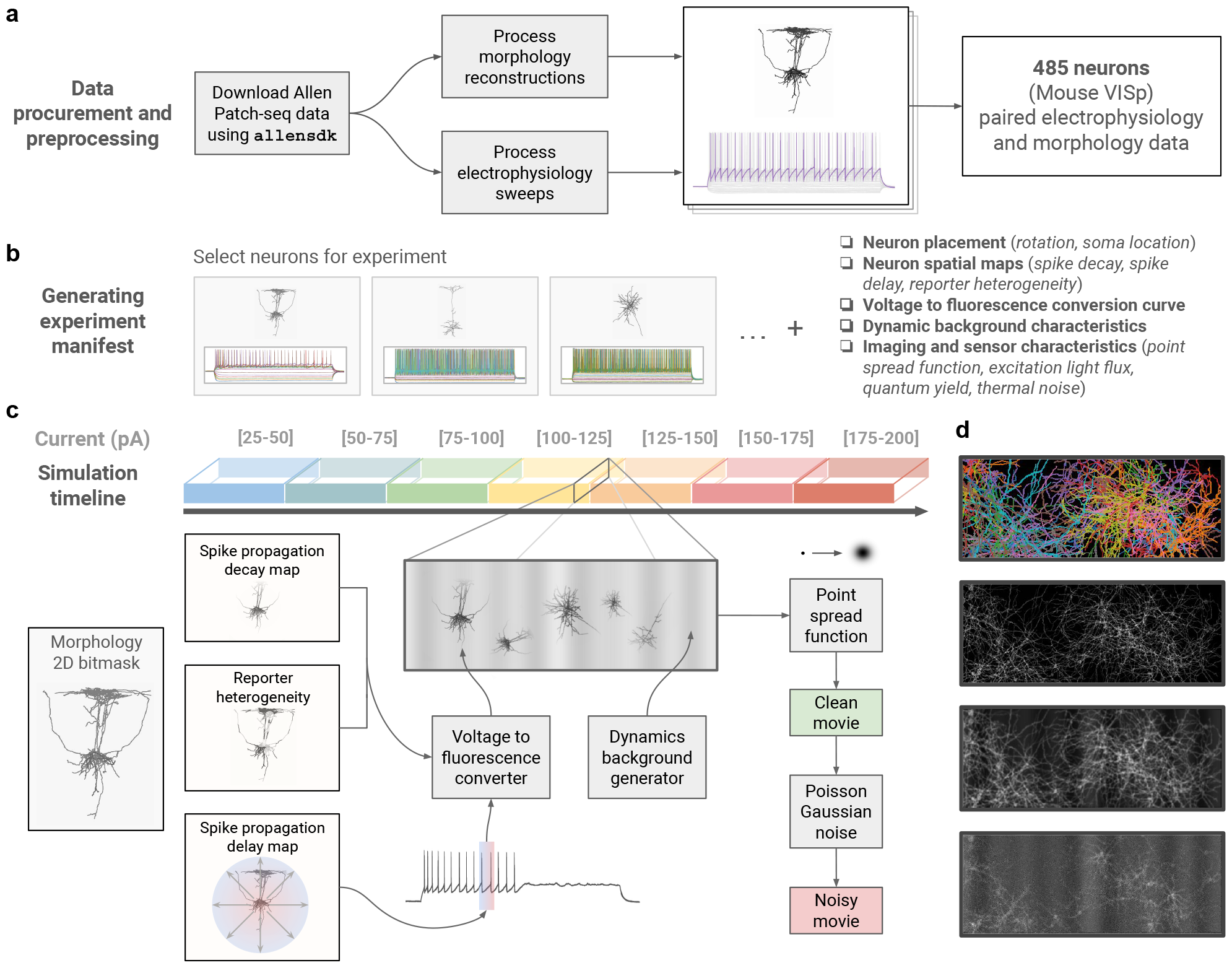
Overview of Optosynth voltage imaging simulation environment. (a) Single-neuron paired morphology and EP data downloaded from Allen Brain Atlas; (b) Generating experiment manifest, including selecting neurons and sweeps for each segment of the experiment, and random sampling and precomputing various simulation accessories; (c) Schematic illustration of the generation process of a movie frame: depending on the position of a pixel on a given neuron, an action potential wavefront propagation delay is read off from the precomputed delay map and is used to select the appropriately delayed timepoint on the EP voltage trace. The voltage value is converted to fluorescence amplitude in combination with the precomputed reporter heterogeneity and spike decay maps. This process is repeated within an efficient vectorized algorithm for all pixels for a given neuron and for all other neurons in the simulation. A background frame is generated and added to the total fluorescence amplitude map generated by the neurons. A point spread function (Gaussian blur) is applied to the total fluorescence map to generate a *clean movie* frame. The application of pixelwise Poisson-Gaussian noise with specified parameters (thermal noise strength, quantum yield) generates a *noisy movie* frame. This process is repeated for each frame in the stimulation segment and for all other segments in the simulation. (d) From top to bottom: (1) neuronal masks juxtaposed in different colors; (2) a simulated frame before the addition of background and PSF; (3) the same frame after the addition of background and PSF; (4) the same frame after the addition of Poisson-Gaussian noise.

We sample *ρ*(**r**) from a Gaussian process with zero mean and an isotropic Gaussian kernel with lengthscale *𝓁*_reporter_. The *f*_clamp_ function linearly rescales the dynamic range of the randomly generated density map to the specified lower and upper limit, 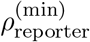 and 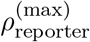, respectively. We randomly place each neuron on the imaging field, together with a random rotation, displacement, and scale factors. To this end, we take the initial binary mask of the *i*’th neuron and apply a general isotropic affine transformation on the mask to obtain the final mask on the imaging field, which we refer to as M^(*i*)^(**r**) *∈* [0, 1]. The precomputed spatial maps and the geometric placements on the imaging field are held constant for all stimulation segments in the experiment, as well as potentially other simulated trials involving the same neurons.

The spatial fluorescence density emanating from a pixel at position **r** at time *t* emanating from the *i*’th neuron, *F* ^(*i*)^(**r**, *t*), is calculated as follows:

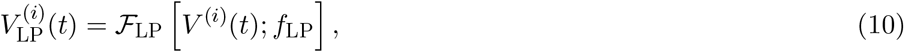

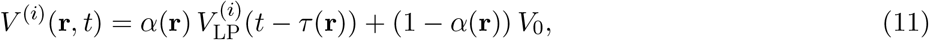

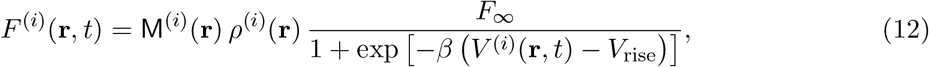

First, we apply a low-pass Fourier filter *ℱ*_LP_ with cutoff frequency *f*_LP_ on the patch-clamp EP recording of *i*’th neuron, *V* ^(*i*)^(*t*), to obtain 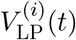. This provision is to simulate the lagged response of one’s choice of fluorescent reporter to voltage transients. To obtain the effective spatial membrane potential of the neuron, *V* ^(*i*)^(**r**, *t*), we linearly admix the resting potential *V*_0_ *≈ −*70 mV and the appropriately time-delayed and low-passed EP recording, see Eq. 11. Finally, the fluorescence density is obtain by converting the spatial membrane potential to fluorescence and multiplying by the precomputed reporter density and neuron mask, see Eq. (12). We use a sigmoid function to convert voltage to fluorescence, and set the sigmoid slope *β ≪* 1 to work mostly in the linear regime. The two parameters *F*_*∞*_ and *V*_rise_ are automatically determined by the user’s specification of two points on the voltage conversion curve, (*V*_1_, *F*_1_) and (*V*_2_, *F*_2_).

#### Simulating the background fluorescence density

Typical voltage imaging recording is often accompanied by a source background fluorescence, including the static autofluorescence from the cell culture, and a dynamic slowly-varying component due to convection currents associated with sample heating by the stimulation laser. We simulate the background noise by first sampling a set of patterns from a zero-mean Gaussian process with anisotropic Gaussian kernels:

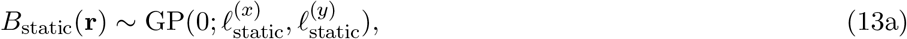

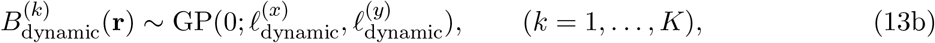

where 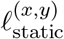 and 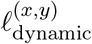 denote the spatial variation lengthscales of static and dynamic background patterns along the *x* and *y* axes, and *K* is the number of dynamic patterns. For each dynamic pattern, we additionally sample a slowly-varying temporal amplitude:

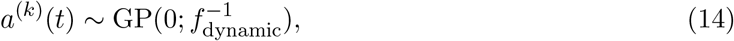

where *f*_dynamic_ is the dynamic background variation frequency. We finally compose the full background as follows:

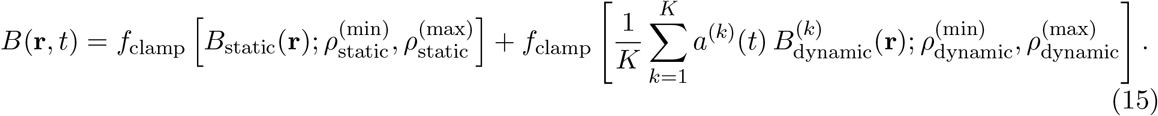

#### Generating the clean and noisy recordings

The total fluorescence density is the sum of fluorescence density associated with the neurons and the background:

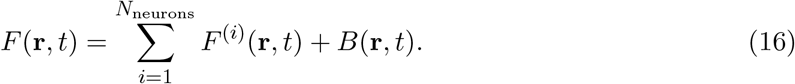

We interpret the fluorescence density *F* (**r**, *t*) as the density of excitable fluorophores per voxel, such that the reporter fluorescence laser converts *F* (**r**, *t*) to an emitted photon count per imaging interval via a conversation factor *Q*. To account for point spreading due to imaging optics, we further convolve *F* (**r**, *t*) with a normalized Gaussian point spread function (PSF) with lengthscale *𝓁*_PSF_ to obtain the clean fluorescence density:

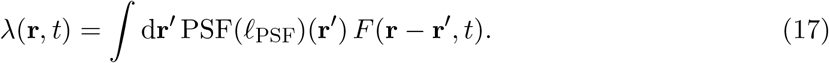

The clean recording is given as:

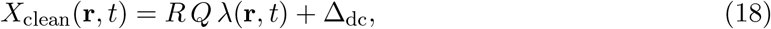

where *R* is the sensor voltage gain per absorbed photon, *Q* is the photon emission rate per excited fluorophore, and Δ_dc_ is a dc offset (characteristic of typical sensor readouts). The noisy recordings are obtained by applying a Poisson-Gaussian noise and quantizing to integer counts:

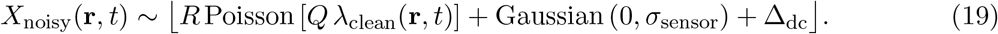

Note that the SNR is implicitly controlled by the two parameters *Q* (number of photons per fluorophore) and *σ*_sensor_ (sensor noise).

To generate multiple simulated recordings from the same neurons, we simply repeat the simulation process multiple times while keeping the experiment manifest constant (including choice of Patch-seq neurons, as well as their geometries and precomputed spatial maps).

### S.6 Optimizing CellMincer network architecture and training schedule using Optosynth-simulated datasets

We performed an extensive hyperparameter exploration to study the role of various modeling choices and identify optimal settings. Simulated data produced by Optosynth was an ideal setting for optimization experiments and ablation studies because of the access to ground truth imaging and its capacity to simulate a range of imaging conditions. The ground truth imaging enabled the direct computation of performance metrics for model evaluation, while the simulation versatility allowed us to test the model on data *imaged* at various SNR without the overhead associated with real-world data collection. Using Optosynth, we generated five datasets under the same neuron density and SNR conditions, three of which were allocated to the training set while the other two were set aside for testing.

Because a complete grid search was not feasible, we chose a baseline configuration that produced a viable model and varied the parameters around this baseline one at a time. Some choices, such as the use of an Adam optimizer, the learning rate scheme, and the number of training iterations, were decided in the baseline model and do not appear in our optimization experiments. We determined variations on each of the other hyperparameters of interest to apply to our baseline model and trained a CellMincer model with each of the resulting configuration variants. These models were subsequently used to denoise both the training and testing datasets, and we computed the distribution of PSNR gain over each frame within the active stimulation periods. The distributions of these PSNR gains and all of our model variants are summarized in Supplementary Fig. S 2.

Our key finding was that the inclusion of global features produced a dominating gain in PSNR. On both training data and unseen test data, CellMincer models incorporating global features exhibited an additional 5 dB gain in PSNR over the baseline model (i.e. a striking 3-fold increase in SNR), in contrast with other architectural variations that yielded 0-1.5 dB over the baseline (Supplementary Fig. S 2a-c). Indeed, we found that the baseline model performance (which did not include the global features) was highly sensitive to changes in the temporal window length (i.e. the denoising context size), with longer windows significantly improving performance. It is evident that both modifications to this baseline model address the limited temporal context it is provided, of which global features is by far the more effective and computationally efficient option. To determine the architectural settings that best synergize with the inclusion of global features, we performed this optimization experiment again with a parallel set of CellMincer variations, all of which included *repeat* global features (Supplementary Fig. S 2d-f). While most of the model performance variation remained consistent with our results in the first iteration, we observed that the performance gain induced by larger temporal contexts plateaued at a window length of 13 frames, comparable in size to our baseline window of 9 frames. This further supports our hypothesis that very long temporal windows can be exploited by a model without global features as a compensatory measure, and by explicitly including global features, we removed the need for long contexts. The result is a network architecture that requires comparatively fewer input frames for denoising, allowing it to denoise datasets in a fraction of the computational time needed by comparable deep learning architectures like DeepCAD.

**Figure S 2:**
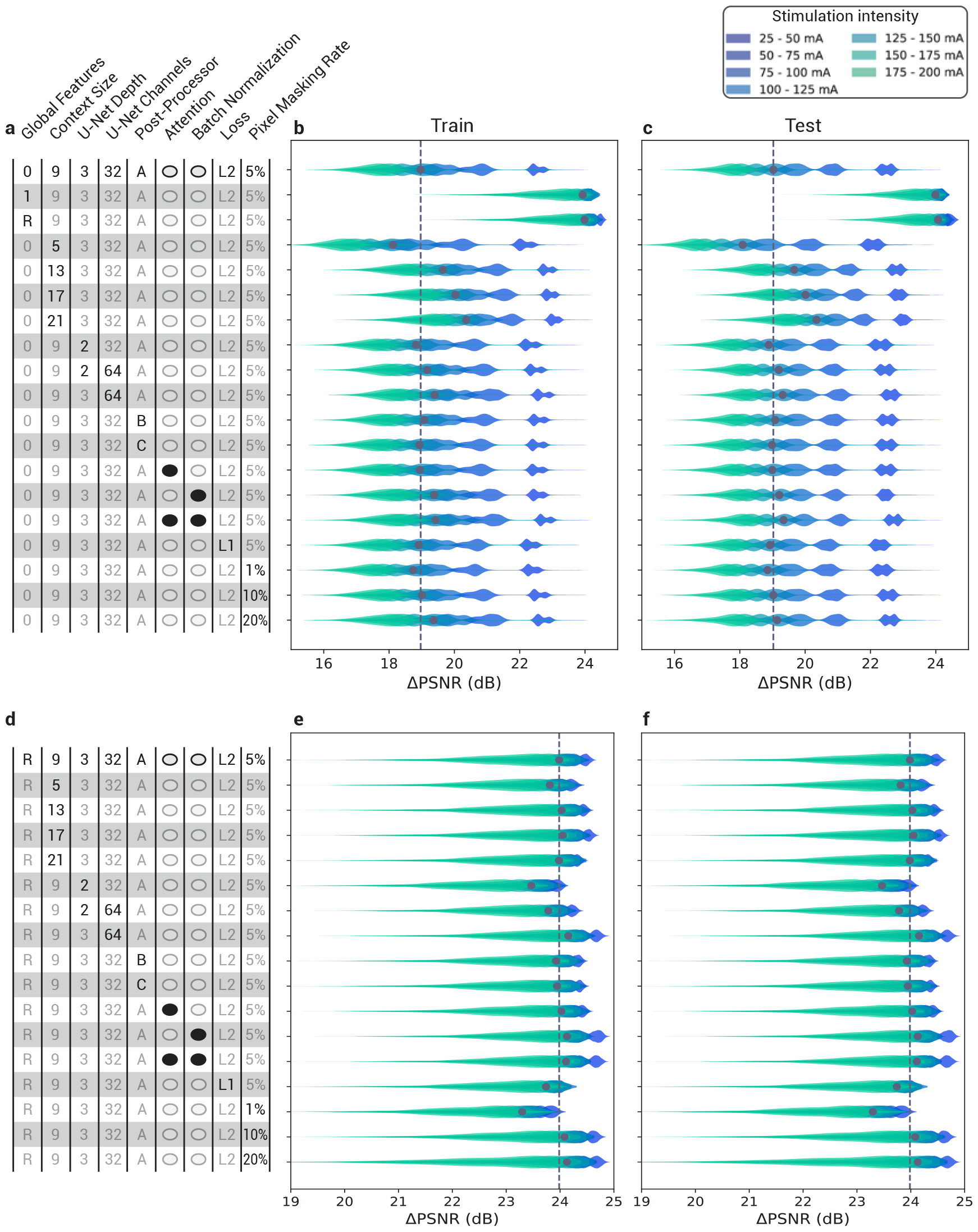
CellMincer hyperparameter settings and their resulting models’ performance on Optosynth data. Each model was evaluated on both its training data (b, e) and unseen test data (c, f). (a)-(c) Initial series of experiments using no global features as a baseline. (d)-(f) Followup iteration of experiments using repeated global features as a baseline. The global features setting determines whether the precomputed global features is not used (0), used to augment the U-Net input only at the beginning (1), or used to augment repeatedly at every contracting path step (R). The included temporal post-processor variants refers to the architecture of the ultimate multilayer perceptron component: *C → C/*2 *→ C* (A), *C → C → C/*2 *→* 1 (B), and *C → C → C →* 1 (C). The architectural variants are ordered in increasing complexity. The pixel masking setting refers to the Bernoulli parameter used to decide whether each pixel is masked, a sampling process repeated for each training iteration. The second set of experiments adjusts the original baseline model to use a conditional U-Net with repeated global features.

### S.7 Metrics for evaluating denoising performance on Optosynth simulated datasets

Peak signal-to-noise ratio (PSNR), measured in decibels, is a metric of similarity between a clean image and its noisy realization and is defined as:

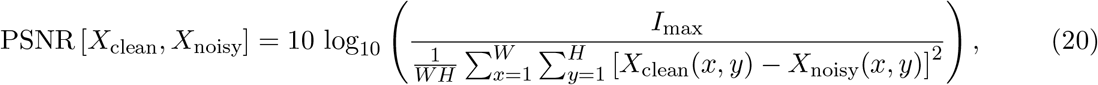

where *I*_max_ is the maximum possible value for the signal intensity. Many of our results are reported in PSNR *gain*, in which we use the PSNR between the raw noisy data and the clean data as a baseline. Reporting the results in terms of PSNR gain is more meaningful and comparable across different settings as it does not depend on *I*_max_. Structural similarity index measure (SSIM) is another metric describing the perceived quality of noisy digital images. We chose to reports on results in terms of PSNR given the flexibility it affords (e.g. the ability to be restricted over arbitrary spatial regions such as foreground or background pixels), as well as its wide adoption in the fluorescence imaging community.

### S.8 Procuring fluorescence intensity traces and aligning to joint electrophysiology data

Our analyses of single-neuron traces are contingent on identifying representative ROIs over which the fluorescent signal is averaged, which is also a common practice in the field. To determine neuronal ROIs, we select a small set of seed pixels that belong to a neuron’s soma and calculate cross-correlations between these pixels and every other pixel in the raw recording. On the resulting cross-correlation map, we choose a manually tuned threshold that captures the soma region and apply the watershed algorithm to add spatial continuity. The resulting ROI is then used to compute traces for the raw recording as well as its denoised counterparts.

Our ROI-extracted fluorescence trace can be interpreted as a noisy affine transformation of the neuron’s electrophysiological activity. In the absence of a calibration dataset, we rely on an optimization-based approach to align the obtained fluorescence traces (in arbitrary units) to the EP data (in mV). We first remove the trend from the imaging trace by median filtering with a moving one-second window (which is short enough to correct for pipette movement but long enough to retain the actual signal). We also removed high-frequency jitter from the patch-clamp EP by applying a Savitzky-Golay cubic filter with a 51-point window. These post-filter signals are more easily imposed over one another following several linear transformations. By matching corresponding peaks in both signals, the intensity trace can be transferred onto the EP timescale. This allows us to evaluate intervals between peaks in absolute time and to downsample the EP signal with interpolation. We then find the affine transformation of our intensity trace that minimizes L2 error with the EP signal, producing an aligned voltage imaging trace. Our subsequent analyses center on evaluating the reconstruction quality of these aligned traces. Refer to Fig. 3b for the results of this alignment method.

### S.9 Metrics for evaluating denoising performance on real voltage imaging with paired EP

#### Quantifying residual noise power using short-time Fourier transform

To compute the spectral noise power of a residual aligned fluorescence trace, we apply a short-time Fourier transform (STFT) parameterized by window length 64 and overlap 48. While voltage imaging, which exhibits a relative downsampling factor of 100, cannot fully capture the underlying EP signal, the peaks of spiking events are particularly high-magnitude points of uncertainty, making them ill-suited for this analysis. To remove spiking events from consideration, we exclude time intervals in which the EP signal’s total spectral power exceeds a chosen threshold. For the remaining time intervals, we convert the spectral powers to noise intensity (dB) and average them in each frequency bin to produce the average noise at that frequency. By repeating this process for each denoised signal and the raw signal, we can compute the reductions in noise intensity, yielding the results shown in Fig. 3d. A typical spectrogram is also shown here in Supplementary Fig. S 3, in which we clearly notice spiking events as and the attenuated high-frequency noise in the inter-spike interval in the CellMincer-denoised results. In contrast, the spectrogram of the raw data is visually akin to that of white noise.

**Figure S 3:**
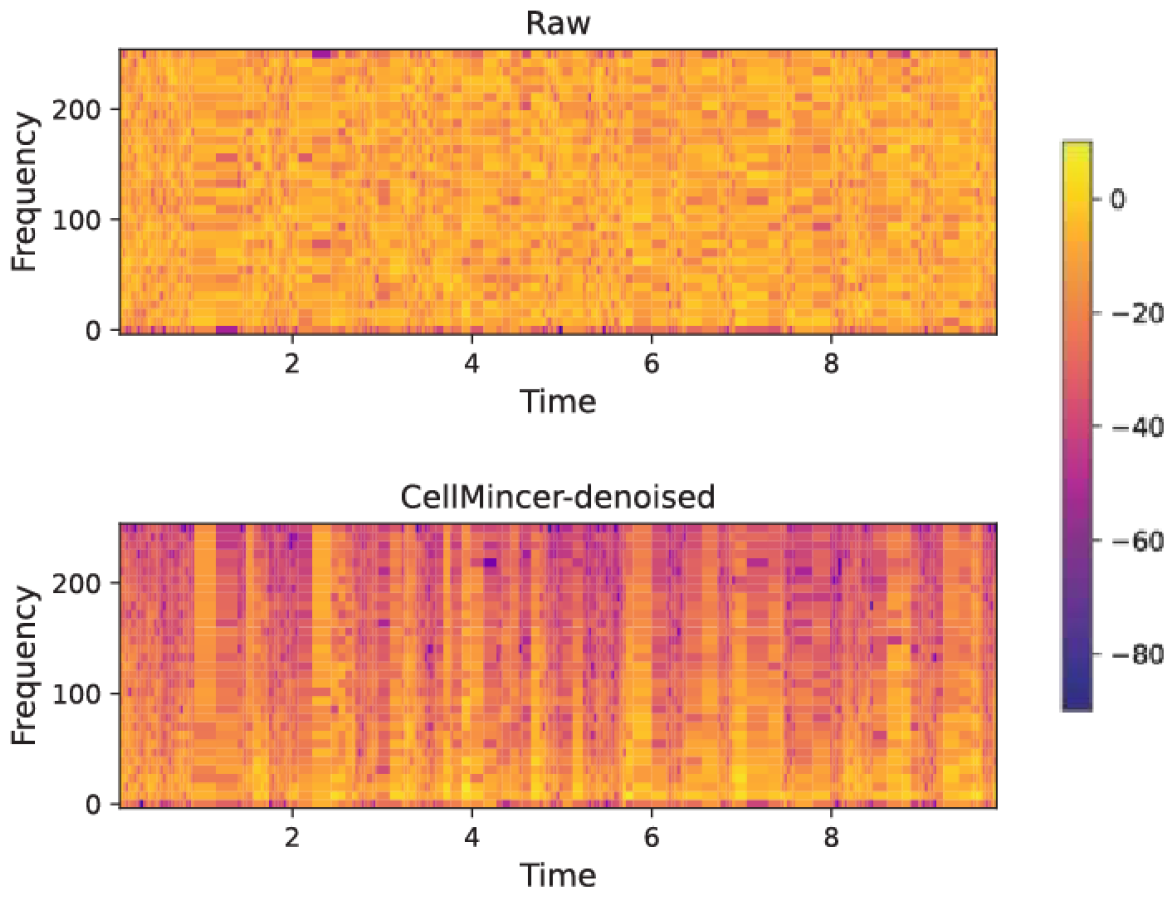
A sample spectral power map (dB) of a residual voltage imaging recording before and after denoising with CellMincer.

#### Prominence-based peak calling

We can formulate the notion of quantifying signal reconstruction quality as a peak-calling problem. We consider peaks in the EP signal and classify them by their prominence, as most spikes exceed 20 mV in prominence while peaks in the subthreshold activity fall below 10 mV. A visualization of prominence as a signal peak feature is shown in Supplementary Fig. S 4. Let *X*_EP_(*t*) and *X*_VI_(*t*) refer to the filtered and aligned traces derived from the patch-clamp EP and the voltage imaging (VI) respectively, and let *S*_*p*_(*X*) be the set of peak time-points in signal *X*(*t*) thresholded above a certain prominence *p*. We express our problem as an evaluation of similarity between *S*_*p*_(*X*_EP_) and *S*_*p*_^*′*^ (*X*_VI_) for *p*^*′*^ *≃ p*. As our set elements comprise points along a continuous time interval, we need to adapt the notions of precision and recall to allow fuzzy matching of nearby peaks. More explicitly, we define:

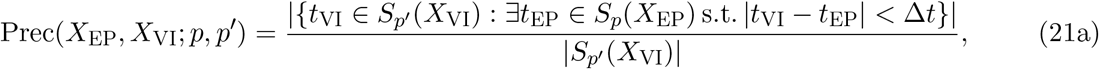

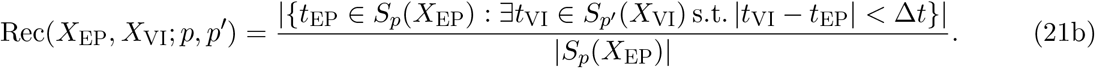

We calculate the *F*_1_-score as usual:

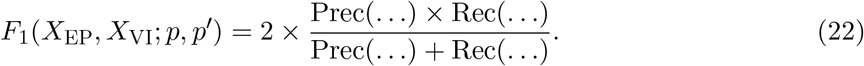

As our definitions suggest, we determine peaks to be correctly called when a corresponding peak occurs within a Δ*t* separation. In our evaluations, we set Δ*t* = 2 ms, corresponding to a one-frame discrepancy in 500 Hz voltage imaging. To introduce tolerance around particular choices of prominence thresholding, we further define:

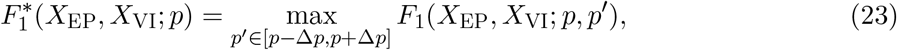

and we set Δ*p* = 0.5 mV. We average this quantity over bins of prominences *p* to produce Fig. 3e, which is the ultimate result of this analysis.

**Figure S 4:**
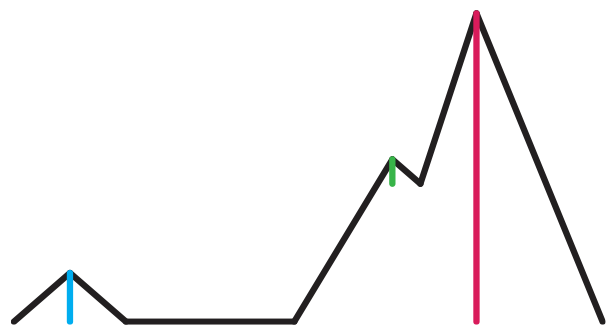
A visualization of the prominence attribute in a simplified signal with three peaks. A moderate prominence threshold would exclude transient peaks produced by noise fluctuations (green), and a larger threshold would exclude subthreshold activity (blue), leaving only the true action potentials (red).

### S.10 Method for segmenting and spike-counting voltage imaging datasets

We extract single-neuron segments from each raw movie and its denoised counterpart using a PCA/ICA-based approach [22]. Similar to CellMincer’s data preprocessing step, our first step is to enhance pixel-pixel correlations by detrending the movie, leaving only the signal component stemming from neural activity. Through experimentation with various signal filters, we found that a rolling circle filter parameterized with width 3.2 frames (6.4 ms) and height 16 (fluorescence a.u.), followed by a threshold filter to collapse values between *±*15 (fluorescence a.u.) to zero, was effective at isolating spiking events in the denoised data. We could not achieve similar success using various combinations of filters on the raw data, so we left those datasets unchanged. Using movie pixels as samples and time frames as features, PCA reduces the dimension of each pixel from *N*_time_ *∼* 10^4^ to *N*_PCA_ = 200. These components correspond to groups of covarying pixels but not necessarily to individual neurons. For this reason, ICA is used to subsequently un-mix the principal components into an independent component set containing our neuron segment candidates. To detect neurons of varying signal strengths, we use a range of parameterizations *N*_ICA_ ∈ {10, 20, 50, 100} to produce the components from which our neuron segments are selected through manual filtering, including a careful study of ICA spatial components together with the associated temporal trace around spike-like events.

Following the extraction of these segments, we compute inner products between a segment mask and each movie frame and apply median filtering with window length of 51 frames to generate its corresponding trace. From these traces, we identify spiking events through manual filtering. This process is aided by a peak-finding algorithm to determine spike candidates and a segmented movie visualization to resolve ambiguous spiking sources. These spikes are tallied across stimulation intensities, and statistical separation between the spiking activity of unperturbed and chronically TTX-treated neurons is computed with a Wilcoxon rank sum test.

It should be noted that through the process of computationally and manually filtering segments, candidates with no discernible activity were excluded from analysis, even though they may have been real neurons which failed to express either the light-gated ion channel or fluorescent reporter. This distinction was considerably more apparent in the denoised movies, potentially causing some neurons to be discarded from the denoised count and included in the raw count despite the overall neuron count favoring the denoised data.

**Figure S 5:**
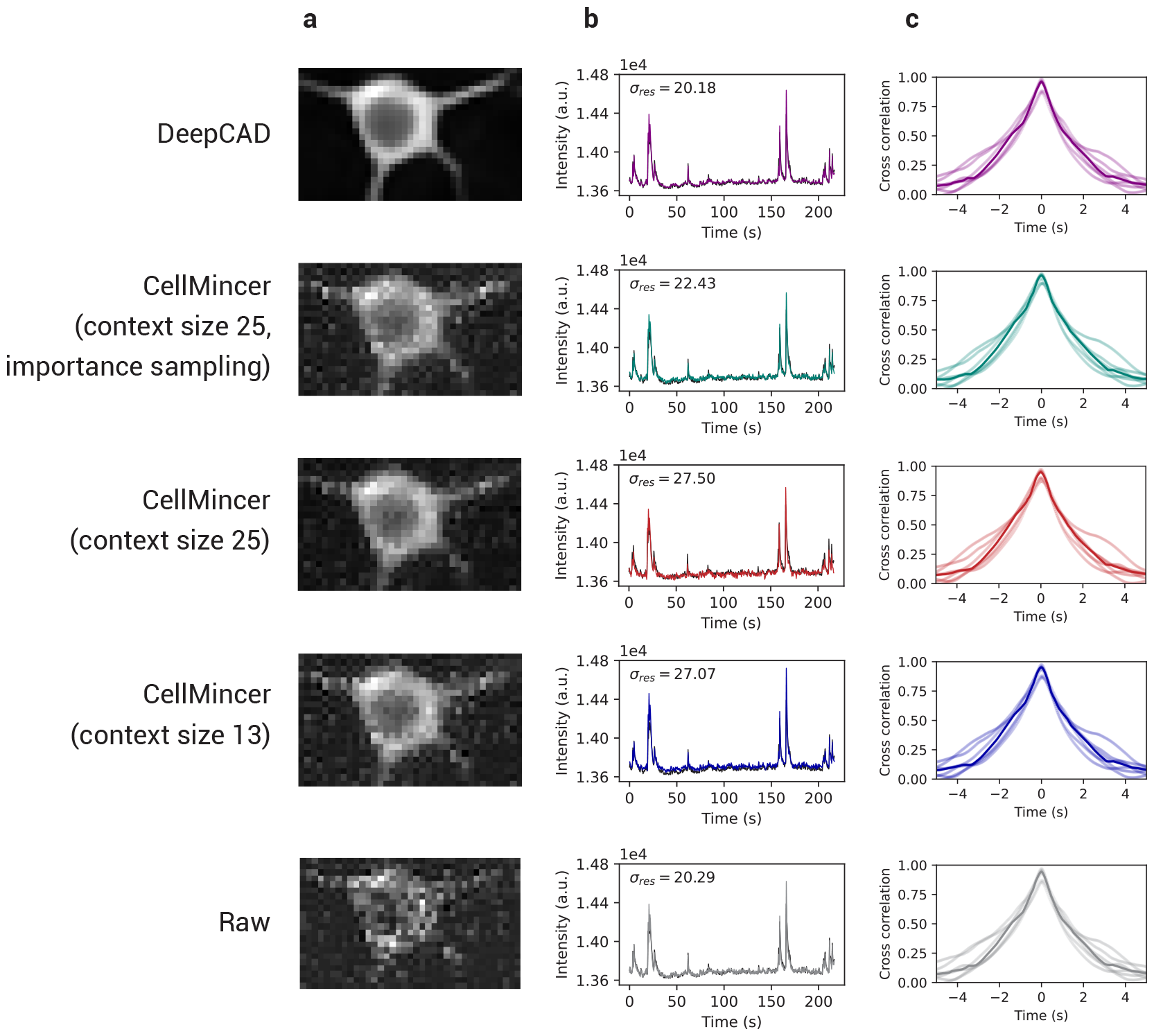
Benchmarking experiments for several configurations of CellMincer and DeepCAD on calcium imaging. In addition to the CellMincer configuration tuned on Optosynth data, we tested two variants which incorporated a significantly longer context size of 25 frames (up from 13 frames), as well as an implementation of importance sampling to increase the frequency of data crops with discernible neural activity in training batches. (a) Sample denoised frame, zoomed in on a select neuron. (b) Averaged intensity trace over sample frame. The standard deviation of the residual signal with respect to the aligned high-SNR trace is labeled. Note that *σ*_res_ was computed over the residual signal following an alignment transformation. This transformation was optimized for the alignment to raw data, which explains the raw signal’s surprisingly low residual variance. (c) Cross-correlations between the denoised trace and high-SNR trace.

### S.11 Limitations of CellMincer vs. DeepCAD on calcium imaging datasets

To first demonstrate an application of the DeepCAD model in its native problem domain, we conducted a limited experiment to compare the efficacy of CellMincer and DeepCAD on calcium imaging. We trained CellMincer and DeepCAD on a training set of seven low-SNR calcium imaging datasets and compared the outputs with their corresponding high-SNR recordings. The results of this experiment, shown in Supplementary Fig. S 5, include multiple iterations of CellMincer configurations that incorporate adaptations to calcium imaging data domain. One adaptation significantly increases the temporal context length to more closely resemble the architecture of DeepCAD, while a second version applies the importance sampling approach used to more efficiently sample active regions of the recording (see Discussion). For the sample neuron presented in Supplementary Fig. S 5a, we found that DeepCAD produced a discernibly cleaner image of the neuron than both CellMincer and the raw data. We then averaged the intensity over each image in column a to produce corresponding single-neuron traces, which we plotted in column b. Overlaid onto these plots is the trace derived from the high-SNR recording aligned to its raw low-SNR counterpart. Ignoring the residual spread of our raw alignment, which is already minimized by construction, we found that while DeepCAD’s trace conforms comparatively well to the high-SNR recording, our importance sampling strategy significantly reduced the performance gap between CellMincer and DeepCAD on calcium imaging. This experiment warrants future research in a more versatile self-supervised training approach that allows denoising both slowly-varying (calcium imaging) and rapidly-varying (voltage imaging) functional imaging modalities.

**Table S 1:**
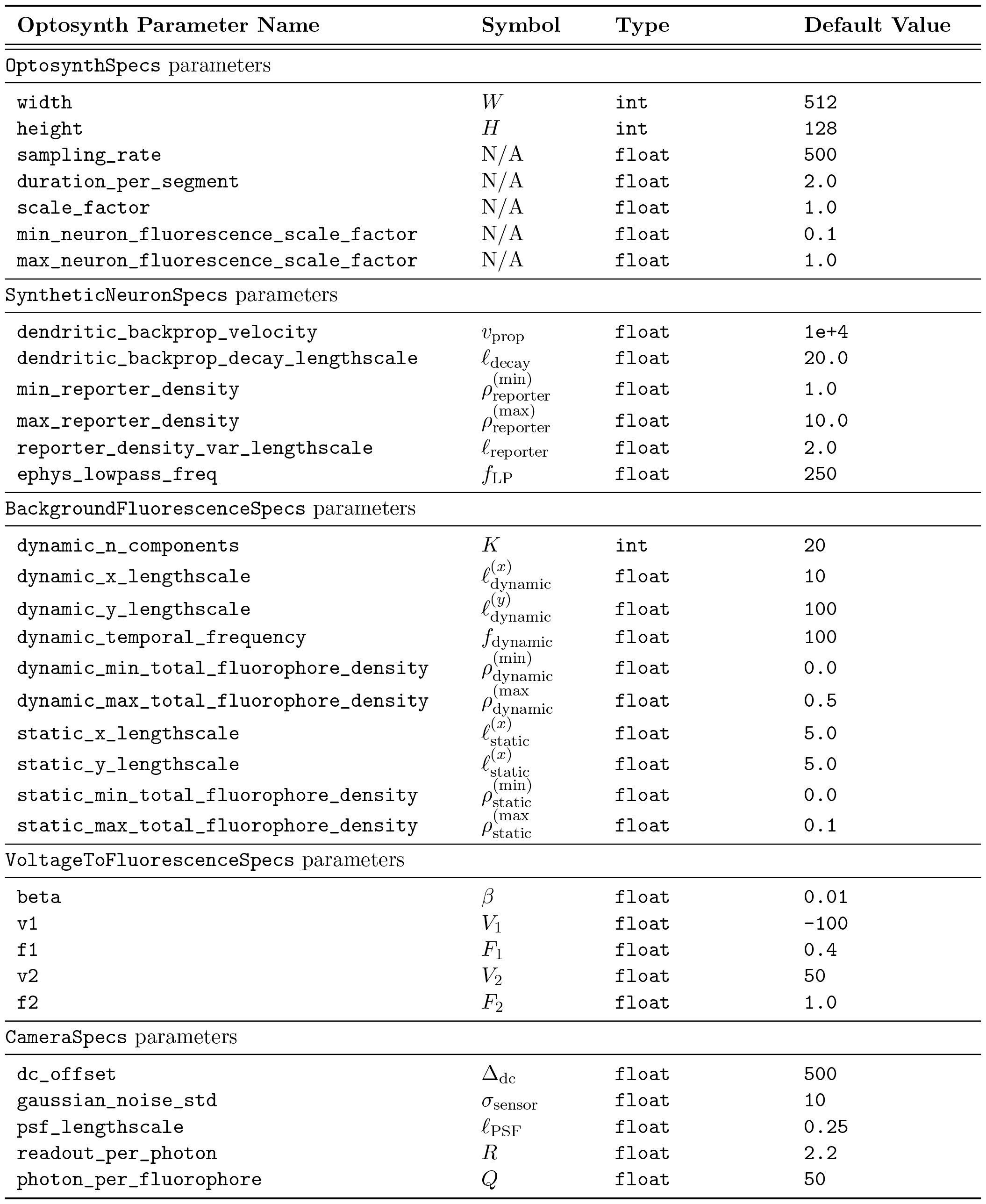
Optosynth parameters and their default values.

| Optosynth Parameter Name | Symbol | Type | Default Value |
| --- | --- | --- | --- |
| OptosynthSpecs parameters |  |  |  |
| width | $W$ | int | 512 |
| height | $H$ | int | 128 |
| sampling_rate | N/A | float | 500 |
| duration_per_segment | N/A | float | 2.0 |
| scale_factor | N/A | float | 1.0 |
| min_neuron_fluorescence_scale_factor | N/A | float | 0.1 |
| max_neuron_fluorescence_scale_factor | N/A | float | 1.0 |
| SyntheticNeuronSpecs parameters |  |  |  |
| dendritic_backprop_velocity | $v_{\text{prop}}$ | float | 1e+4 |
| dendritic_backprop_decay_lengthscale | $\ell_{\text{decay}}$ | float | 20.0 |
| min_reporter_density | $\rho_{\text{reporter}}^{(\text{min})}$ | float | 1.0 |
| max_reporter_density | $\rho_{\text{reporter}}^{(\text{max})}$ | float | 10.0 |
| reporter_density_var_lengthscale | $\ell_{\text{reporter}}$ | float | 2.0 |
| ephys_lowpass_freq | $f_{\text{LP}}$ | float | 250 |
| BackgroundFluorescenceSpecs parameters |  |  |  |
| dynamic_n_components | $K$ | int | 20 |
| dynamic_x_lengthscale | $\ell_{\text{dynamic}}^{(x)}$ | float | 10 |
| dynamic_y_lengthscale | $\ell_{\text{dynamic}}^{(y)}$ | float | 100 |
| dynamic_temporal_frequency | $f_{\text{dynamic}}$ | float | 100 |
| dynamic_min_total_fluorophore_density | $\rho_{\text{dynamic}}^{(\text{min})}$ | float | 0.0 |
| dynamic_max_total_fluorophore_density | $\rho_{\text{dynamic}}^{(\text{max})}$ | float | 0.5 |
| static_x_lengthscale | $\ell_{\text{static}}^{(x)}$ | float | 5.0 |
| static_y_lengthscale | $\ell_{\text{static}}^{(y)}$ | float | 5.0 |
| static_min_total_fluorophore_density | $\rho_{\text{static}}^{(\text{min})}$ | float | 0.0 |
| static_max_total_fluorophore_density | $\rho_{\text{static}}^{(\text{max})}$ | float | 0.1 |
| VoltageToFluorescenceSpecs parameters |  |  |  |
| beta | $\beta$ | float | 0.01 |
| v1 | $V_1$ | float | -100 |
| f1 | $F_1$ | float | 0.4 |
| v2 | $V_2$ | float | 50 |
| f2 | $F_2$ | float | 1.0 |
| CameraSpecs parameters |  |  |  |
| dc_offset | $\Delta_{\text{dc}}$ | float | 500 |
| gaussian_noise_std | $\sigma_{\text{sensor}}$ | float | 10 |
| psf_lengthscale | $\ell_{\text{PSF}}$ | float | 0.25 |
| readout_per_photon | $R$ | float | 2.2 |
| photon_per_fluorophore | $Q$ | float | 50 |

